# Coupling cell communication and optogenetics: Implementation of a light-inducible intercellular system in yeast

**DOI:** 10.1101/2022.06.25.497625

**Authors:** Vicente Rojas, Luis F. Larrondo

## Abstract

Cell communication is a widespread mechanism in biology, allowing the transmission of information about environmental conditions. In order to understand how cell communication modulates relevant biological processes such as survival, division, differentiation or apoptosis, different synthetic systems based on chemical induction have been successfully developed. In this work, we coupled cell communication and optogenetics in the budding yeast *Saccharomyces cerevisiae*. Our approach is based on two strains connected by the light-dependent production of α-factor pheromone in one cell type, which induces gene expression in the other type. After the individual characterization of the different variants of both strains, the optogenetic intercellular system was evaluated by combining the cells under contrasting illumination conditions. Using luciferase as a reporter gene, specific co-cultures at 1:1 ratio displayed activation of the response upon constant blue-light, which was not observed for the same cell mixtures grown in darkness. Then, the system was assessed at several dark/blue-light transitions, where the response level varies depending on the moment in which illumination was delivered. Furthermore, we observed that the amplitude of response can be tuned by modifying the initial ratio between both strains. Finally, the two-population system showed higher fold-inductions in comparison with autonomous strains. Altogether, these results demonstrated that external light information is propagated through a diffusible signaling molecule to modulate gene expression in a synthetic system, which will pave the road for studies allowing optogenetic control of population-level dynamics.

**Graphical Abstract:** 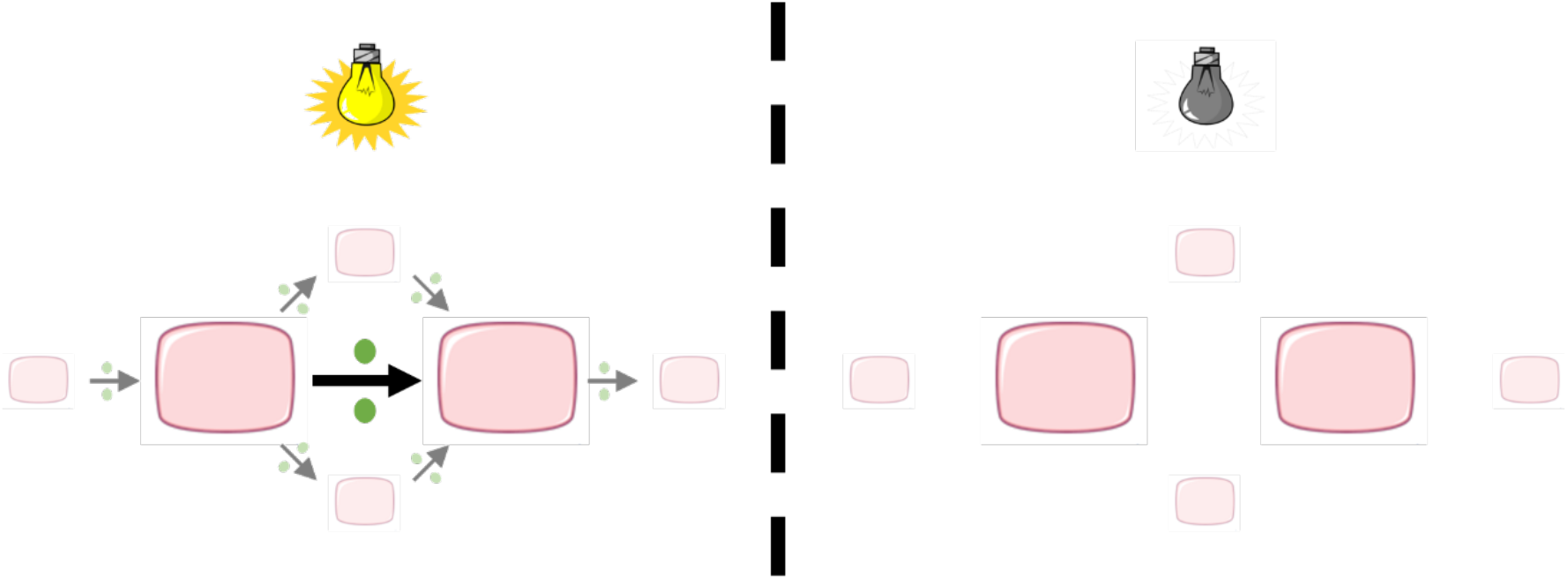

## INTRODUCTION

Communication is an essential process in living organisms, allowing the efficient transmission of valuable information ^1,2^. In cells, different communication mechanisms have been selected throughout evolution, that can be classified into two main types. The first relies on direct contact between cells ^3^, whereas the second one depends on the secretion of signaling molecules ^4,5^. Notably, the latter has been used to implement synthetic systems with the aim of understanding how cell communication regulates biological processes in response to environmental conditions. These systems require gene circuits that lead to the inducible production of signaling molecules, establishing cell communication in a predictable manner ^6–8^. To date, there are several examples of synthetic intercellular systems that have been implemented in prokaryotic and eukaryotic platforms, allowing the study of different behaviors at the population level ^9–14^. However, most of them are induced by chemicals. Despite its high activation levels, this induction strategy has limitations such as irreversibility, low spatiotemporal resolution and toxicity ^15^. In recent years, optogenetics has emerged as a promising alternative to replace chemicals ^16^. This technology utilizes natural or engineered photoreceptors that sense light of well-defined wavelengths, improving the spatiotemporal resolution and reducing unwanted effects of traditional inducers ^17–20^. But although light works as a reversible cue that overcomes several problems, synthetic approaches coupling cell communication and optogenetics have been seldom explored so far ^21,22^.

The budding yeast *Saccharomyces cerevisiae* has served a pivotal role as a model organism to dissect eukaryotic biological processes ^23^. This unicellular fungus has tractable genetics, facilitating the design of synthetic genetic circuits by the addition of non-native building blocks or the modification of endogenous molecular pathways ^24,25^. Moreover, yeast has two additional features that make it an ideal host to implement an optogenetic intercellular system. First, *S. cerevisiae* is not able to perceive light as it lacks functional photoreceptors ^26^. Nevertheless, plenty of research has reported the successful implementation of optogenetic switches in yeast, where light acts as an orthogonal input ^27,28^. On the other hand, *S. cerevisiae*’s life cycle involves a communication mechanism based on small diffusible pheromones, that serve as signaling molecules ^29,30^. The binding of these pheromones to specific G protein-coupled receptors in the cell surface triggers the mating process, which involves the activation of a mitogen-activated protein kinase (MAPK) cascade leading to chemotropic growth by large-scale changes in gene expression, cell division cycle and morphology ^31–33^. As *S. cerevisiae* lacks motility mechanisms, this pheromone-dependent polarized extension allows haploid yeast cells to get close, make contact and fuse to form a diploid zygote ^34,35^.

Several studies have manipulated the native mating process at the molecular level to control the secretion of pheromones and the biological outcomes of the pathway upon specific stimuli ^36–38^. In that context, *S. cerevisiae* has already been used as chassis to implement synthetic intercellular approaches, using pheromone signaling to establish artificial communication by rewiring its original components. For instance, chemicals such as galactose, estradiol and NaCl have been used to express α-factor in a set of engineered yeast cells. Different combinations of them with a reporter strain led to the generation of simple Boolean circuits ^39^. Similarly, more complex logic gates have been implemented in a yeast consortium using progesterone, aldosterone and dexamethasone as exogenous inducers to activate the production of the mating pheromone ^40^. In order to avoid potential intracellular cross-talk, yeast cells have also been engineered to develop synthetic systems where communication depends on plant signaling molecules such as auxins ^41,42^. Importantly, synthetic intercellular approaches are not restricted to using a single population as occurs in autonomous systems, in which each cell is containing all the components to establish communication and its concomitant response. In fact, the increasing complexity of synthetic circuits has caused the generation of a new type of system, where the components to perform specific biological processes are compartmentalized in two or more non-isogenic populations ^43–45^. Thus, the different strains can be interconnected as a modular assembly line by cell communication, avoiding metabolic burden, reducing stochastic noise, and increasing genomic stability ^46–49^.

Considering all the aspects mentioned above, we developed a two-population system where light sensing and the expression of a luciferase gene reporter are physically separated, but they are readily connected by cell communication based on a small diffusible molecule such as the α-factor pheromone. The response of the system is quite versatile since gene expression levels depend on the specific cell mixture, the illumination condition, and the initial ratio between both strains. In relation to autonomous versions of the circuit, specific symmetric co-cultures reached lower levels of response but a better performance in terms of the fold-induction between blue-light and dark conditions. In summary, the results suggest that optogenetic intercellular approaches could constitute a powerful tool to synchronize phenotypes in yeast, improving the current understanding of dynamics in the transmission of environmental regulatory signals.

## RESULTS AND DISCUSSION

FUN-LOV, an optogenetic switch we recently described by exploiting the blue-light dependent interaction between WC-1 and VVD photoreceptors from the filamentous fungus *Neurospora crassa*, has enabled a fine temporal resolution of transcriptional control yielding a broad range of expression in yeast ^50^. In this work, we used this switch to implement a light-inducible intercellular system based on two yeast strains (Figure 1). The first strain (called OS, for <u>O</u>ptogenetic <u>S</u>ender) harbors FUN-LOV and produces α-factor upon blue-light by activating the transcription of the *MFα1* pheromone-encoding gene. Then, this secreted molecule binds to Ste2 surface receptors of the second strain (called R, for <u>R</u>eceiver), activating the mating pathway that leads to the expression of a luciferase reporter gene under the control of the *FUS1* pheromone-responsive promoter. As detailed below, we first proceeded to individually characterize the OS- and R-strains, and then, the effects of combining them.

**Figure 1.**
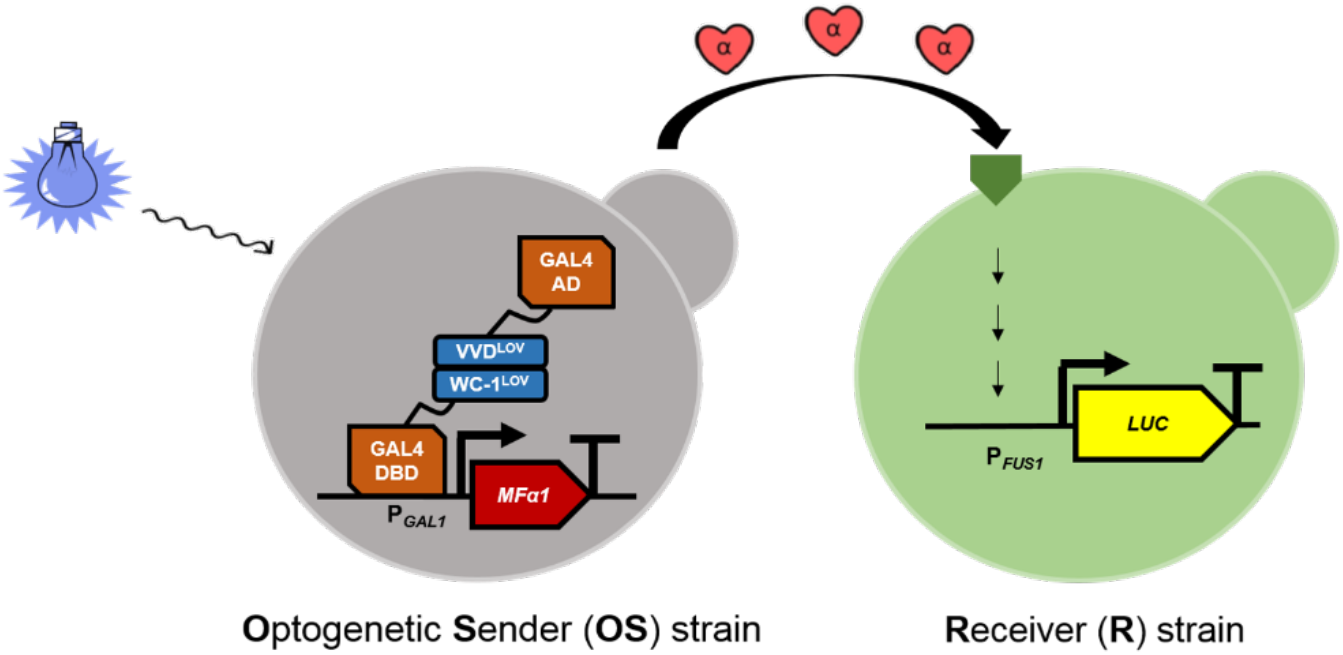
Molecular design of the optogenetic intercellular system. Under blue-light conditions, the FUN-LOV switch activates the secretion of α-factor pheromone by OS-strains. This signaling molecule triggers the mating pathway in R-strains, inducing the expression of luciferase inserted at the *FUS1* locus as a transcriptional reporter.

### R-strains respond to exogenous pheromone

We evaluated the response of R-strains to increasing concentrations of commercial α-factor. It is worth noticing that we generated three variants of R-strain, by inserting the reporter construct into different genetic backgrounds. In this way, R1 was generated by placing luciferase under the control of the *FUS1* promoter in BY4741. This strain required as minimum as 5 µM of exogenous pheromone to induce a strong normalized response (Figure 2). Although the expression of Ste2 pheromone receptor is induced in presence of the pheromone ^51^, the response did not increase at higher concentrations such as 100 µM. In that context, *MATa* yeast strains - like R1 - secrete Bar1 protease as a form to discriminate different pheromone gradients by cleaving α-factor into two inactive fragments ^52,53^. In order to obtain a hypersensitive phenotype, R2 strain was generated by integrating the luciferase construct at the *FUS1* locus in the genome of a *bar1*Δ mutant. As expected, R2 was capable to respond to lower concentrations of supplemented α-factor, such as 1 µM (Figure 2). However, the response peaks remained stable despite adding higher α-factor concentrations to the cultures. On the other hand, a crucial feature of the mating pathway is the synchronization of haploid cells at the G1 stage of the cell cycle to allow fusion prior to DNA replication ^54^. To do this, the MAPK cascade activates the Far1 protein, leading to the inhibition of the Cdc28 cyclin-dependent kinase (CDK), which is one of the main regulators of yeast cell division cycle ^55^. As a result, the mitotic division is temporally blocked. In fact, the OD_600_ curves of R1 and R2 were differentially affected by exogenous pheromone (Figure S1). To avoid these growth alterations in response to α-factor, we used a *bar1*Δ*far1*Δ double mutant to generate R3 by replacing the *FUS1* ORF with the reporter construct. This strain maintained the hypersensitivity of the transcriptional response observed for R2 (Figure 2), but the OD_600_ curves overlapped with the control in the absence of the signaling molecule (Figure S1). Here, the amplitude of the normalized luminescence continued to be unaltered when higher pheromone concentrations are tested, suggesting a saturation or inactivation of the pathway that leads to reporter expression. Besides Bar1 production, *S. cerevisiae* has other mechanisms of α-factor desensitization, including endocytosis of Ste2 receptors ^56^, reconstitution of the G protein that initiates the intracellular signal transduction by Sst2-mediated GTP hydrolysis ^57^ and action of Msg5, Ptp2 or Ptp3 phosphatases to inhibit Fus3 terminal kinase ^58^. In this way, yeast prevents hyperactivation of the MAPK cascade, which leads to stress and cell death ^59^. For that reason, negative regulators of the mating pathway are ideal targets to generate mutants that allow expanding the repertoire of R-strains in the future. Likewise, it would be interesting to evaluate if the maximum response of R-strains increases by using a multi-copy reporter construct.

**Figure 2.**
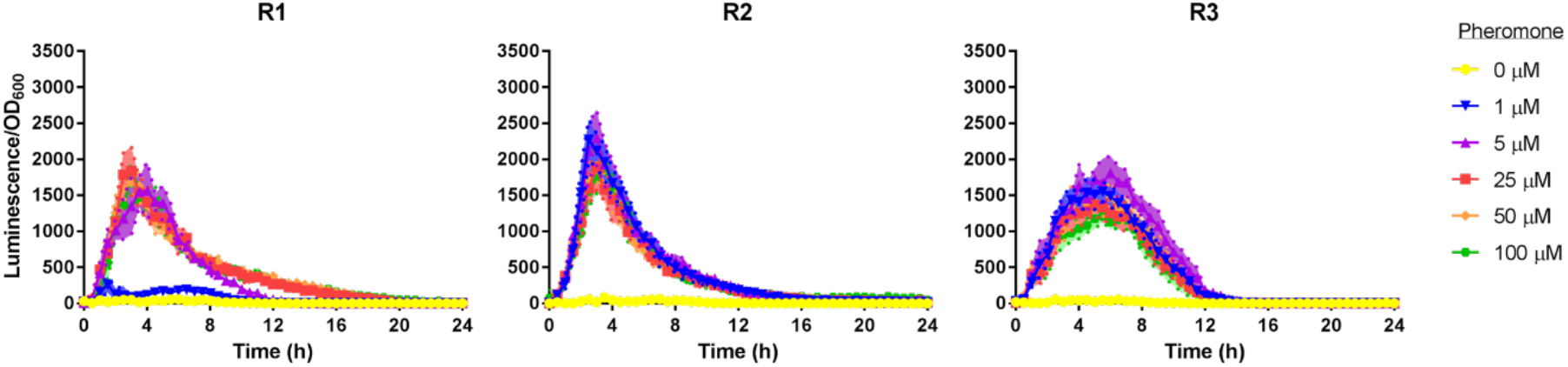
The R-strains depict different profiles of transcriptional response upon pheromone addition. Commercial α-factor was exogenously added in different concentrations (1, 5, 25, 50 and 100 µM) and reporter expression (luminescence normalized by OD_600_) was measured in R1, R2 and R3 cells for 24 h. In all panels, R-strains were evaluated in the absence of pheromone (0 µM) as negative controls. The average normalized response of six biological replicates is shown, with the standard deviation represented as a region with soft tone.

Predictably, no bioluminescence in response to α-factor was detected when yeast cells do not bear luciferase at the *FUS1* locus (Figure S2). In contrast, the temporal growth inhibition of these strains was still observable in the OD_600_ curves, according to their particular genotype, as seen for the R-strains (Figure S2). However, these results showed some discrepancies compared to the halo assays. For instance, inoculation of pheromone on uniform lawns containing R1 cells led to almost imperceptible zones of inhibition only at 50 and 100 µM (Figure S3), unlike what was observed in liquid media. On the other hand, halo assays in R2 indicated a clear dose-dependent growth inhibition (Figure S3), proving to be more informative than the abrupt alteration of OD_600_ curves in the presence of α-factor in liquid cultures. As expected, R3 was not affected by the application of pheromone drops (Figure S3), confirming the results observed in the micro-cultivation experiments. The reason for these differences may be that pheromone diffusion and its interaction with R-strains differ between both culture conditions. Therefore, these results showed that R-strains can display different profiles of transcriptional induction and variable growth alteration upon pheromone addition.

### OS-strains produce functional α-factor in response to light

Naturally, *MATa* cells synthesize the a-factor lipopeptide pheromone, but no α-factor. Conversely, *MATα* cells synthesize the α-factor peptide pheromone, but no a-factor ^29^. Despite that, we generated a genetic construct that allows the inducible secretion of α-factor in *MATa* strains. In that context, *S. cerevisiae*’s genome possesses two pheromone-encoding genes (*MFα1* and *MFα2*) associated with α-factor production. Although both genes are considered paralogs, it has been reported that *MFα1* expression is more active and its polypeptides generate twice the amount of pheromone molecules after processing, contributing to the majority of total α-factor produced by *MATα* cells in basal state ^60^. In order to obtain higher levels of α-factor by optogenetic induction in the OS-strains, we chose *MFα1* to assemble the inducible plasmid.

Some studies have shown that the secretion of α-factor by yeast strains can be directly quantified by approaches based on mass spectrometry ^61^ and ELISA ^62^. However, in this work, the optogenetic production of pheromone by OS-strains was evaluated by its ability to trigger a biological response such as induction of gene expression. After these yeast cells were grown in DD and LL conditions, the supernatants were collected and used to evaluate their functionality in R-strains. In this case, we also generated three variants of OS-strains by co-transforming the FUN-LOV components and the inducible *MFα1* construct in different genetic backgrounds. Thus, OS1 is carrying the three recombinant plasmids in BY4741, and its supernatants were not able to induce any response in R-strains, regardless of the illumination conditions used to incubate the yeast cultures (Figure 3). Following the previous logic, OS2 and OS3 were obtained using the *bar1*Δ and *bar1*Δ*far1*Δ mutants, respectively. In contrast to OS1, supernatant from OS2 cells grown in LL induced a strong normalized response during the first 4 h of cell growth only in the hypersensitive R-strains, which was not observed when we used the supernatant obtained in DD (Figure 3). Similarly, OS3 supernatant collected from LL also induced a transcriptional response in R2 and R3, displaying higher maximum levels followed by a sharp decrease (Figure 3). Nevertheless, both supernatants obtained in LL did not produce a significant response in R1, suggesting that the expression of *BAR1* - by any strain - is decisive to modulate the levels of functional pheromone. Interestingly, the OS2 and OS3 supernatants collected in LL negatively affected the growth curves of the R2 strain (Figure S4). Further analysis of the OD_600_ data showed a mild decrease in growth parameters such as rate and efficiency (Figure S5). In contrast to these results, OS-supernatants drops were incapable of generating growth inhibition assessed by halo assay (Figure S6).

**Figure 3.**
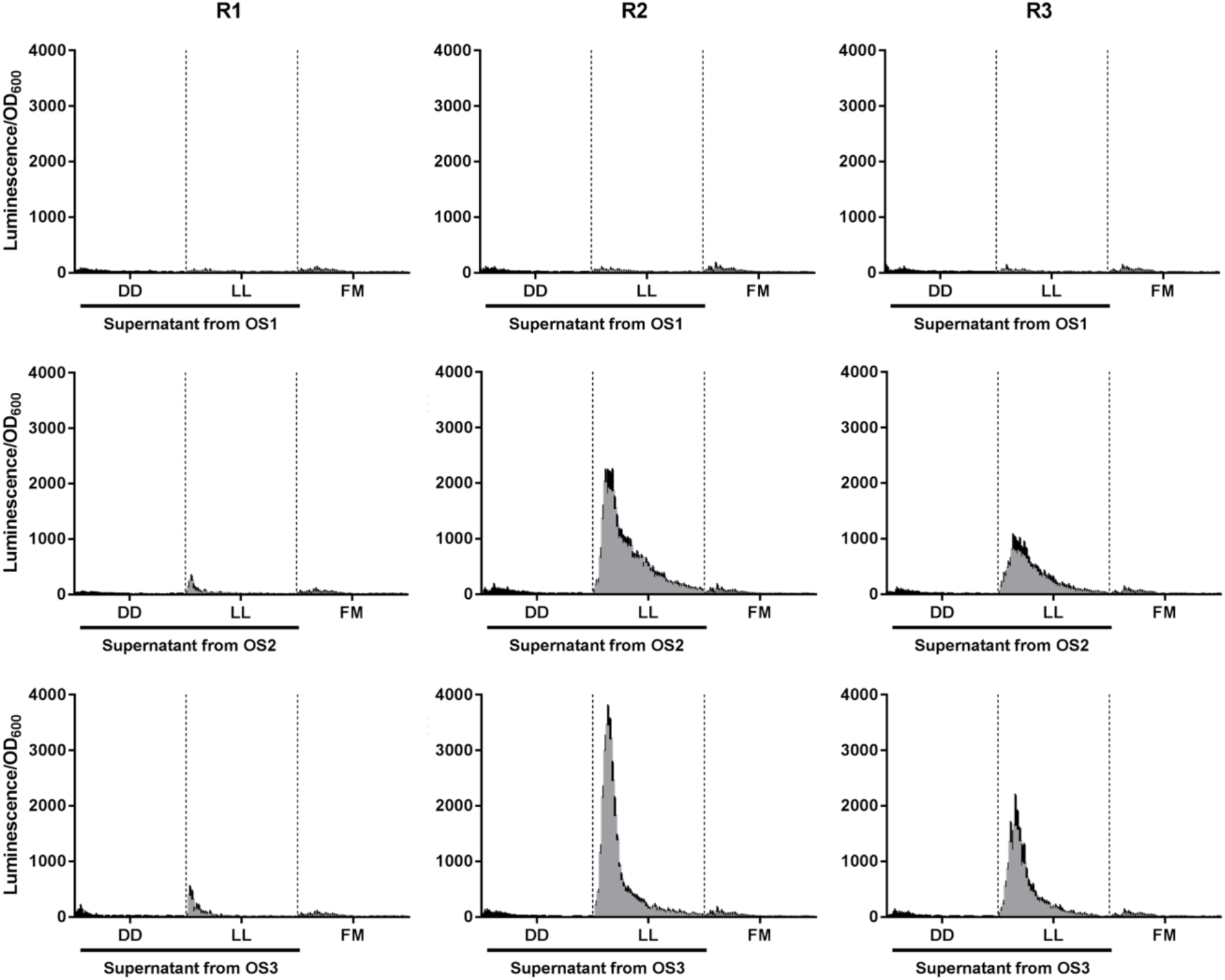
The OS-strains carrying a multi-copy inducible *MFα1* plasmid produce functional pheromone upon constant light. The supernatants of OS1, OS2 and OS3 cultures were collected from contrasting illumination conditions (DD and LL) and reporter expression (luminescence normalized by OD_600_) was measured in R1, R2 and R3 cells for 24 h. In all panels, the R-strains were evaluated in fresh media (FM) as negative controls. The average normalized response of six biological replicates is shown, with the standard deviation represented as a black bar.

Following the same methodology used for generating OS-strains, it would be possible to include in *MATα* cells the optogenetic secretion of a-factor. Although the latter may display solubility issues by its lipophilic nature, synthetic pheromone-dependent bidirectional communication could be implemented in yeast, resembling genetic circuits based on cross-feeding ^63^ or quorum sensing autoinducers ^64,65^. In addition, new OS-strains might be generated by controlling *MFα2* gene expression to provide different response levels and kinetics of the R-strains. Expanding the repertoire of OS-strains can also be carried out by using FUN-LOV variants to increase the level of production of α-factor by light or improve the LL/DD fold-induction ^66^.

To verify if the transcriptional response and growth alteration depended on the optogenetic production of α-factor, we generated OS_control_ strains. The latter maintains the FUN-LOV plasmids, but an empty pRS426 vector replaces the inducible *MFα1* construct. Supernatants collected from OS_control_ strains, grown in DD or LL, did not induce luciferase expression in R-strains (Figure S7). Then, we evaluated the relevance of copy number in the optogenetic induction of the pheromone-encoding gene. In this way, the replacement of the *MFα1* endogenous promoter was carried out to obtain new OS-strains based on a chromosomal inducible construct. Surprisingly, the supernatants of OS4, OS5 and OS6 did not activate the normalized response in R-strains (Figure S8). Altogether, these results indicated that LL condition allows the production of α-factor in OS-strains at enough concentration to activate luciferase transcription in R-strains, as long as Bar1 protease is not secreted and *MFα1* is expressed from a multi-copy vector.

### Light information is propagated by cell communication in a two-population system

After the individual characterization of OS- and R-strains, the optogenetic intercellular system was evaluated by mixing the variants in nine symmetric combinations under DD and BL conditions. Only certain co-cultures at 1:1 ratio and exposed to BL showed activation of the normalized response during the first 10 h, which did not occur for the same cell mixtures grown in DD (Figure 4). Specifically, co-cultures involving R2 reached peaks between 3.5 and 4 h. On the other hand, the kinetics of cell mixtures involving R3 were slower, exhibiting the greatest values between 9 and 9.5 h. Interestingly, these maximum levels were similar to those ones obtained in BL for direct optogenetic control of a chromosomal luciferase using the FUN-LOV switch ^50^. Although blue-light is the stimulus that triggers the split genetic circuit that leads to luciferase induction in the R-strains, the response of the system was surprisingly higher when examining co-cultures going through an illumination transition such as DL4h:20h, compared to ones grown in BL (Figure 4). Here, the maximum levels are distributed over a narrow range of time due to the activation peaks were observed between 5.5 and 6 h. For both conditions, the response progressively decayed after the peak, returning to its basal level at 15 h. Again, there was not a significant transcriptional response when – at least – one of the involved cells bears the *BAR1* gene, confirming the results observed for R-strains evaluated with OS-supernatants. Thereby, the production of this protease neutralizes the presence of the secreted α-factor pheromone, rendering the two-population system unable to establish productive communication. In that context, drops from OS2 and OS3 cultures caused the appearance of inhibition halos in R2 strain only when the plates were grown in LL conditions (Figure S9). As expected, no transcriptional response was observed when we tested co-cultures between OS_control_- and R-strains (Figure S10). In the same way, luciferase expression was not induced when OS, OS_control_ and R strains were evaluated as monocultures (Figure S11). Therefore, these results indicated that non-isogenic yeast strains can interact by cell communication using light as an external controller.

**Figure 4.**
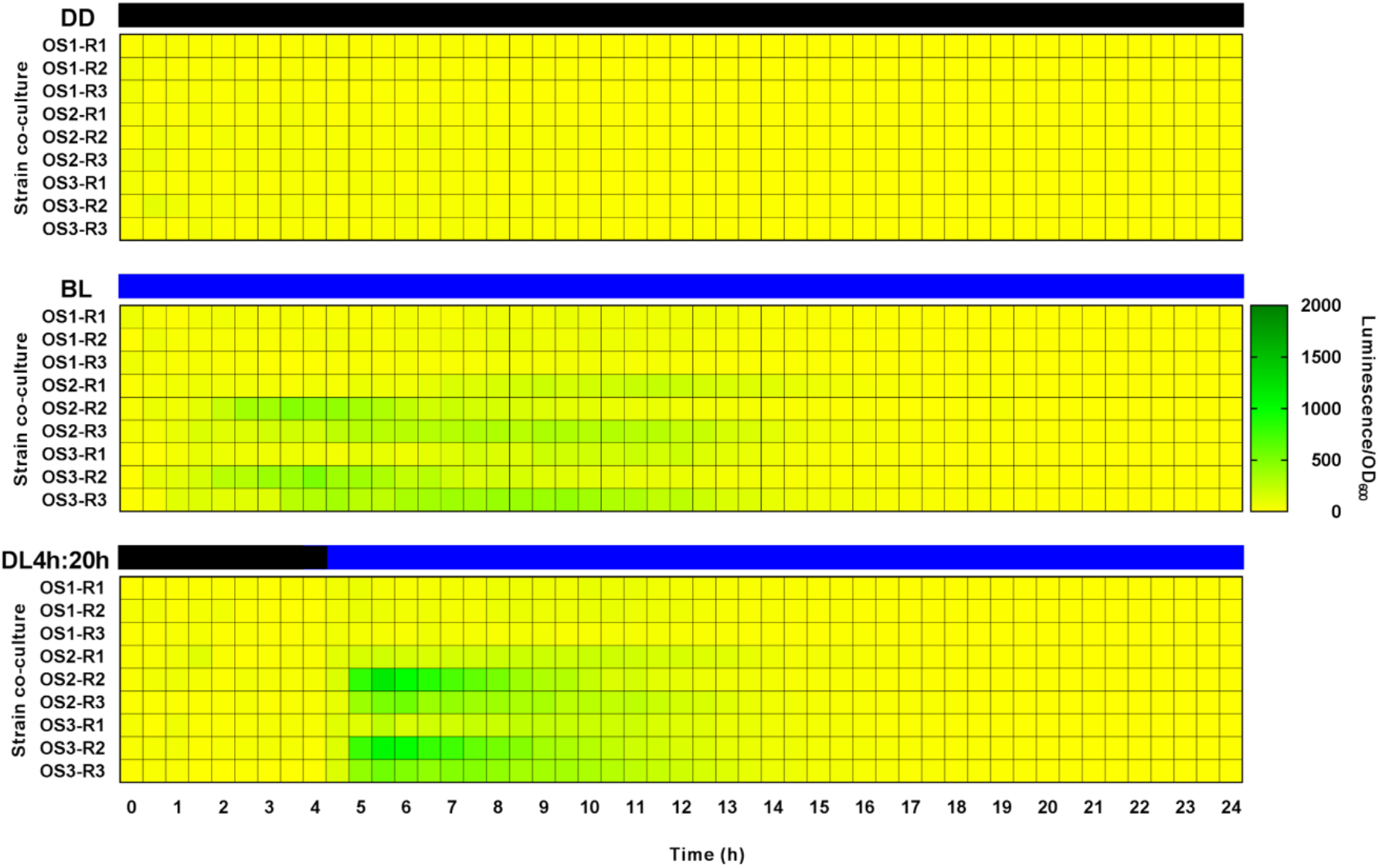
The combination of OS- and R-strains leads to productive light-dependent cell communication. The cells were combined at OS-R 1:1 ratio, and then exposed to DD, BL and DL4h:20h to measure reporter expression (luminescence normalized by OD_600_) for 24 h. The average normalized response of six biological replicates is shown in the heat-maps.

Since the highest response was obtained in DL4h:20h, we evaluated the system under different dark/blue-light transitions by changing the moment at which the LED lamp is turned on. In comparison to DL4h:20h, the system performance was similar at early transitions such as DL1h:23h, DL2h:22h, and DL6h:18h (Figure S12A), with the peaks occurring between 1.5 and 6 h after the blue-light was applied. On the other hand, the yeast co-cultures displayed drastically reduced levels of reporter gene at late transitions such as DL8h:16h, DL10h:14h, and DL12h:12h (Figure S12B). However, although we changed the illumination conditions of the experiments, OS- and R-strains continued to be unable to generate a luminescent response if they were not combined (Figure S13). By using a novel approach based on dynamic fluorescent reporters, the activity of *FUS1* promoter has been detected between 15 and 30 min after α-factor addition ^67^. For that reason, *FUS1* was classified as an early pheromone-responsive gene. Thus, and considering that the activation of the response in our system depends on blue-light administration of OS-R co-cultures, it is not unexpected that the normalized luminescence levels abruptly dropped beyond 8 h of cell growth. In order to obtain a considerable transcriptional response during a posterior window of time, late pheromone-responsive promoters could be used, such as *KAR3* ^67^. The activity of this promoter has been detected between 30 and 50 min after pheromone exposure and its respective gene encodes for a protein related to karyogamy, one of the last steps of yeast mating. In the same context, other variants of the two-population system might be obtained by testing other pheromone-responsive promoters that have been validated in the evaluation of specific processes of interest, including *FIG1* ^68^, *AGA1* ^69^ or an optimized version of *FUS1* ^70^.

Furthermore, we measured the response of the optogenetic intercellular system by evaluating different OS-R ratios. To do this, we changed the inoculation of each type of strain without altering the final volume (10 µL), making comparable the response between symmetric and asymmetric co-cultures. Importantly, the response increased when the initial percentage of R-strains trebled in relation to OS-strains (Figure 5A). In the same way, the normalized luminescence remained stable or decreased in the inverse situation, that is, when the initial volume of OS-strains trebled regarding R-strains (Figure 5B). As seen when the timing of dark/blue-light transitions was modified, these differences in the maximum transcriptional levels were mainly observed in cell mixtures involving OS2 and OS3 with hypersensitive R-strains. However, the temporality of response peaks was maintained in comparison with symmetric co-cultures. Remarkably, a low amount of OS-strains was enough to produce a strong response in R-strains, opening the possibility to implement the system for industrial purposes ^71^. Besides its utility as a host to dissect different cellular phenomena, *S. cerevisiae* is an organism widely used to produce diverse compounds of interest ^72,73^. To increase the yield of these processes, yeast cells are subjected to fermentation in bioreactors, in which cell cultures reach high densities ^74^. Here, chemical induction continues to be the best way to control gene expression, because light is not sufficiently effective to activate all cells in the bioreactor vessel. However, the two-population system would provide a solution to this problem since it is required that only a minimal fraction of cells (OS-strain) senses light to transmit the external input by the action of α-factor, activating gene expression throughout the rest of the culture (R-strain). Importantly, the optogenetic response of specific OS-R mixtures should involve cell cycle arrest at the G1 stage. In that context, it has been suggested that synchronous cultures are useful in yeast biotechnology to optimize the synthesis of high-value metabolites by fermentation ^75^. For instance, the induction of the mating pathway has allowed decoupling cell growth from the production of para-hydroxybenzoic acid ^76,77^.

**Figure 5.**
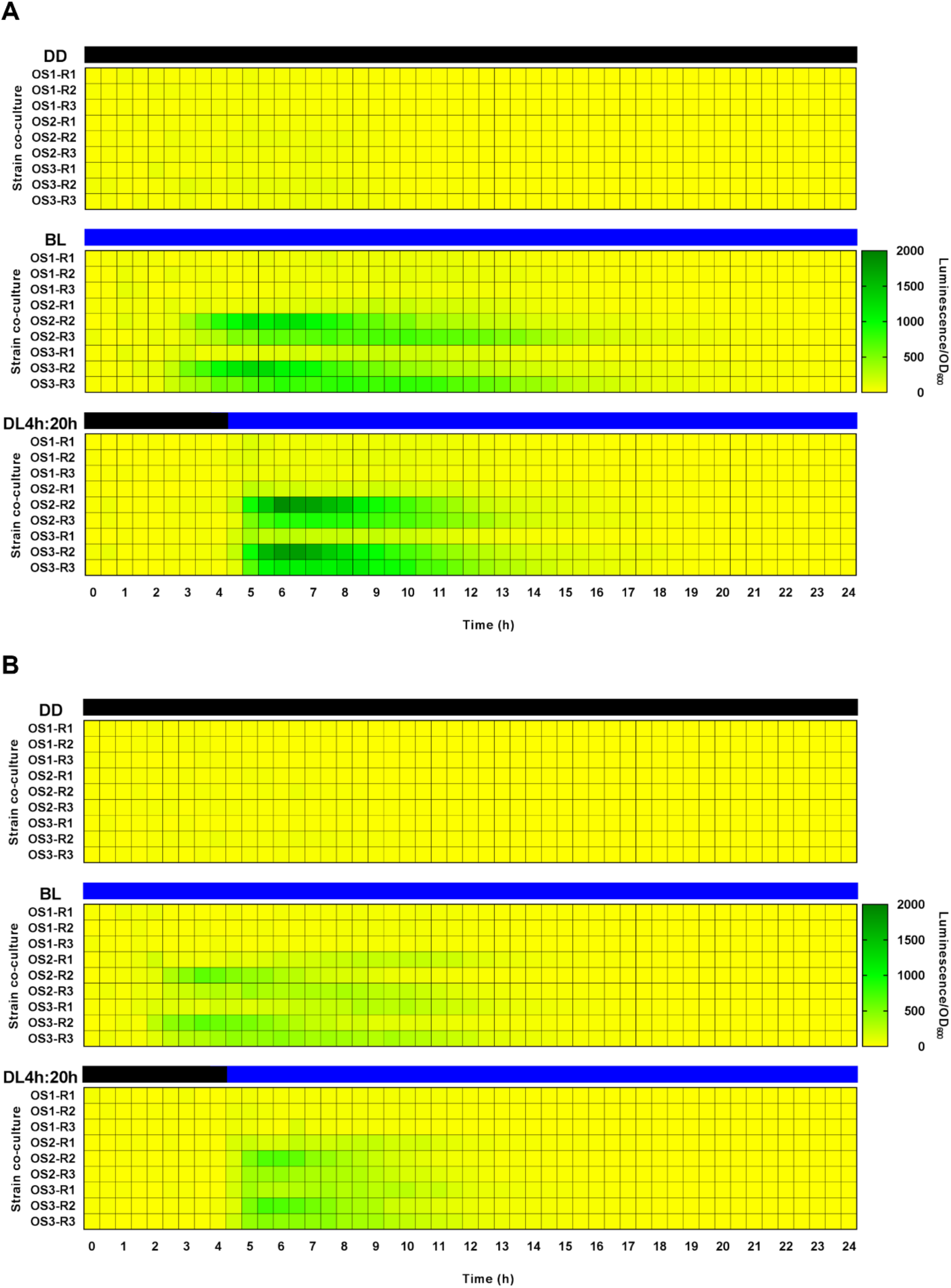
The amplitude of the response depends on the initial ratio between OS- and R-strains. The cells were combined at OS-R 1:3 (A) or 3:1 (B) ratios, and then exposed to DD, BL and DL4h:20h to measure reporter expression (luminescence normalized by OD_600_) for 24 h. The average normalized response of six biological replicates is shown in the heat-maps.

Finally, autonomous (A) versions of our synthetic circuit were generated, where each cell is carrying all the building blocks related to light perception, pheromone production and reporter expression (Figure 6A). Thus, each cell of A-strains has the capacity to induce luciferase expression in response to optogenetically-produced pheromone. In contrast, the *FUS1* promoter activity is limited by the initial percentage of R-strain in the two-population system. For instance, only 50% of all cells could produce bioluminescence in OS-R co-cultures at 1:1 ratio. Compared with the symmetric co-culture approach, A-strains showed slightly higher levels in response to BL and DL4h:20h, but also higher basal activity in DD (Figure 6B). In this way, the fold-inductions of A-strains were lower than ones related to certain OS-R mixtures at 1:1 ratio (Figure 6C), indicating that the separation of biological activities in two different strains improves the performance by filtering the transcriptional noise ^48^. This result was similar or even better whether A-strains are compared with the asymmetric OS-R co-cultures (Figure S14). As expected, no response was observed when we tested A_control_-strains (Figure S15).

**Figure 6.**
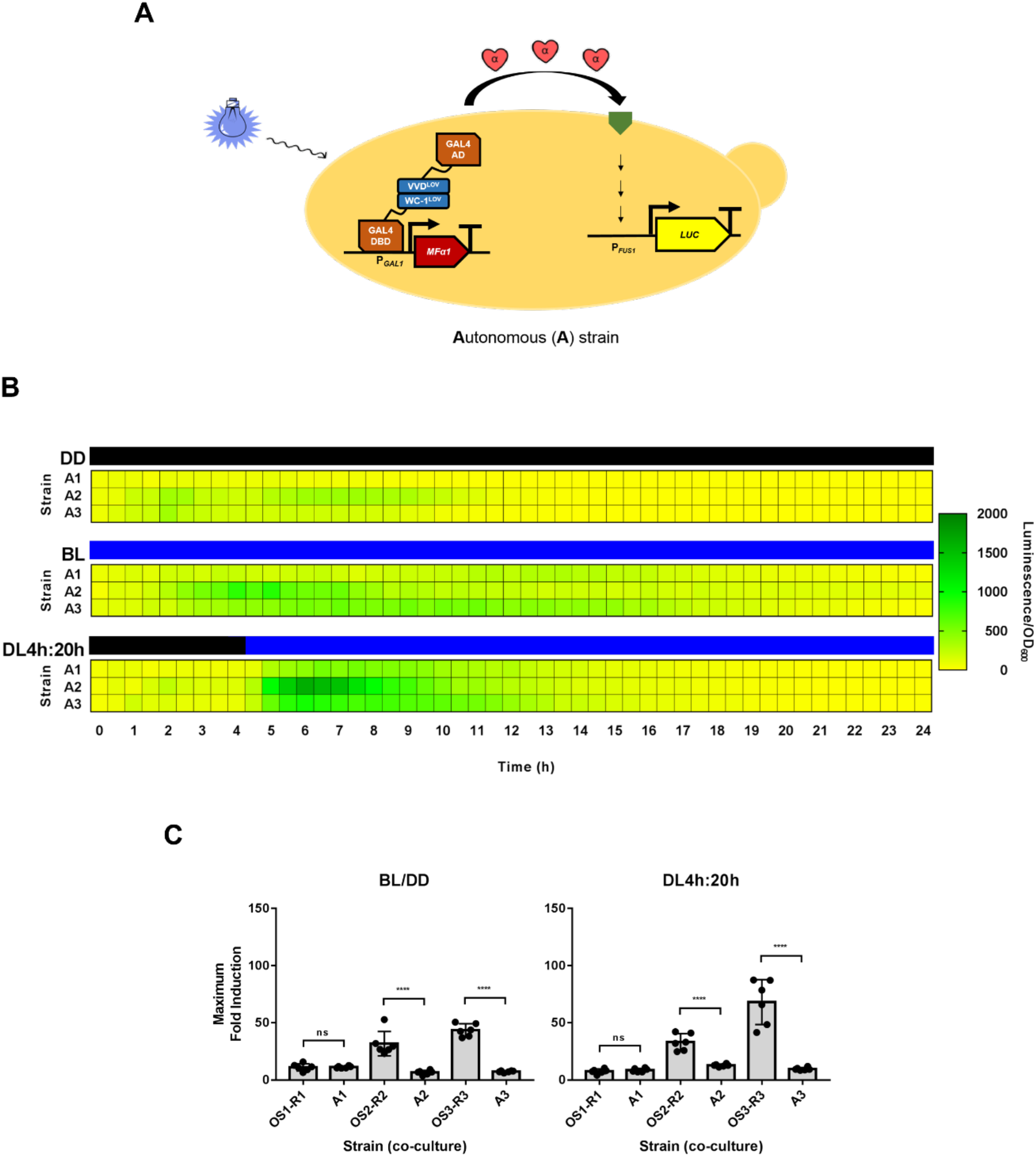
The autonomous strains display a worse performance regarding symmetric OS-R co-cultures. A. Molecular design of A-strains. B. The cells were exposed to DD, BL and DL4h:20h to measure reporter expression (luminescence normalized by OD_600_) for 24 h. The average normalized response of six biological replicates is shown in the heat-maps. C. Comparative of maximum fold-induction between A-strains and the optogenetic intercellular system at OS-R 1:1 ratio for two illumination conditions (****P <0,0001).

The design of the two-population system pretends that pheromone produced by OS-strains triggers the mating pathway in R-strains (Figure 1). Nevertheless, this scheme omits certain biological activities that would be occurring in the co-culture experiments. One of them is that part of the secreted α-factor by light probably binds to surface pheromone receptors of the same OS-cells, that lack the luciferase reporter gene. Thus, OS-strains are likely acting as sponges, reducing the effective concentration of signaling molecules that can be used by R-strains to generate a measurable response. In that context, two strategies are being implemented to avoid this loss of α-factor. The first option is the evaluation of co-cultures between A- and R-strains. Similar experiments involving mixtures between a “secrete-and-sense” strain and an “only-sense” strain have shown that the response resembles the quorum sensing behavior ^78,79^, where a threshold concentration of molecules is reached while cell density increases, generating a coordinated response at population level mediated by autocrine and paracrine communication. The second option is the generation of OS-strains that also carry a *STE2* deletion. Furthermore, it is important noticing that specific cell co-cultures may involve nutrient competition between the respective strains. For instance, when OS2 (whose mitotic division is affected by pheromone) is combined with R3 (its cell cycle remains unaltered in presence of α-factor), the latter might overgrow the former regardless of the initial ratio between both strains. The same situation is possible in the OS2/R3 mixture. Albeit these concepts are not addressed in this work, the optogenetic intercellular system seems to be useful to assess ecological paradigms, including the appearance of cheaters ^80^.

Future work also aims to obtain more intricate versions of our optogenetic intercellular system. For instance, it would be attractive to evaluate the effect of adding an intermediate strain able to amplify or attenuate the response in R strains, resembling coherent and incoherent feedforward designs. In fact, a three-population circuit was already developed in yeast by exploiting the pheromone signaling ^68^. Using engineered *MATa* strains, the system acts like a chain reaction that propagates the information by the secreted pheromone. However, the *MFα1* gene is constitutively expressed by the first strain. Thus, although the signal is intensified by the second strain to activate the response on the third one, the overall system lacks a user-controlled inducer. In the same way, and considering that the OS-R co-cultures are based on intra-species communication ^81^, the two-component system could be expanded by adding other organisms such as *E. coli* or mammalian cells. Inter-species communication systems, involving *S. cerevisiae* strains, have already been reported using volatile acetaldehyde as a signaling molecule ^82^. Also, the α-factor has been successfully secreted by a modified strain of the fission yeast *Schizosaccharomyces pombe* ^83^. This heterologous expression of the pheromone was validated by cell cycle arrest, morphological changes and induction of mating genes in *S. cerevisiae MATa* cells, confirming the possibility to implement our system with different organisms. These types of synthetic circuits may enable the formation of consortia that contributes improving the knowledge about the propagation of external stimuli at population level.

## METHODS

### Yeast strain and culture conditions

*S. cerevisiae* strain BY4741 (*MATa; his3Δ1; leu2Δ0; met15Δ0; ura3Δ0*), was used as the genetic background for yeast transformation ^84^. This strain was maintained in YDPA medium (2% glucose, 2% peptone, 1% yeast extract and 2% agar) at 30ºC. Co-transformants carrying plasmids with auxotrophic markers were maintained in synthetic media (0.67% yeast nitrogen base without amino acids, 2% glucose, 0.2% amino acids drop-out mix and 2% agar) at 30ºC.

### Generation of yeast mutant and reporter strains

The one-step PCR deletion by recombination protocol ^85^ was followed to generate gene mutants of the mating pathway. Specifically, the kanamycin (*KanMx*) antibiotic resistance cassette was amplified by PCR using a Phusion Flash high-fidelity PCR master mix (Thermo Scientific, USA) and 70-nt primers for direct homologous recombination on the *BAR1* locus, allowing the swapping of the endogenous ORF region in the BY4741 strain. Then, the nourseothricin (*NatMx*) antibiotic resistance cassette was amplified and transformed in the BY4741 *bar1Δ* strain to generate the allelic swapping on the *FAR1* locus. The same procedure was used to generate the pheromone-responsive strains. A previously described luciferase reporter construct ^86^ was amplified by PCR using a Phusion Flash high-fidelity PCR master mix (Thermo Scientific, USA) and 70-nt primers for targeted recombination on the *FUS1* locus, allowing the swapping of the endogenous ORF region in the BY4741, *bar1Δ* and *bar1Δfar1Δ* strains. The chromosomal DNA of all strains was extracted using the Wizard® Genomic DNA Purification Kit (Promega, USA) and the integration of antibiotic resistance cassettes or reporter genetic constructs in the genome was confirmed by PCR under standard conditions using the GoTaq Green Master Mix (Promega, USA). The strains used and generated herein are shown in Table S1. Primers used for the generation of strains and their checking are listed in Table S2.

### Generation of recombinant genetic constructs

We used the original FUN-LOV system ^50^ to control the expression of an α-factor pheromone encoding gene (*MFα1*) by light. This inducible genetic construct was generated using yeast recombinational cloning ^87^. Succinctly, the two DNA fragments (*GAL1* promoter and *MFα1* ORF plus its endogenous terminator) were amplified using a Phusion Flash High-Fidelity PCR Master Mix (Thermo Scientific, USA), employing 50-nt oligonucleotides that allow the cloning into pRS426 vector. The presence and orientation of DNA fragments were confirmed by colony PCR using GoTaq Green Master Mix (Promega, USA), whereas the sequence of the genetic construct was checked by Sanger sequencing reactions (Macrogen Inc., Korea). The plasmids used and generated herein are shown in Table S3. The primers used for plasmid assembly are listed in Table S4.

### Evaluation of R-strains using commercial pheromone

The reporter gene coding sequence (*LUC*) was inserted downstream of the endogenous *FUS1* promoter, permitting luciferase expression in response to α-factor pheromone. After that, the reporter strains were co-transformed with empty pRS423, pRS425 and pRS426 vectors, obtaining the Receiver (R) strains. The R-strains were assayed using different concentrations of commercial α-factor pheromone (1, 5, 25, 50 and 100 µM) and the luminescence levels and optical density at 600 nm (OD_600_) of the cell cultures were simultaneously measured over time using a Cytation 3 microplate reader (BioTek, USA). Briefly, yeast strains were grown overnight in a 96-well plate with 200 µL of SC medium at 30ºC. Thereafter, 10 µL of these cultures were used to inoculate a new 96-well plate containing 190 µL of fresh SC media supplemented with luciferin at 1 mM final concentration. The OD_600_ and the luminescence were acquired at 30ºC every 30 min for 24 h, running a protocol with 30 sec of shaking before data acquisition. The kinetic curves were performed with six biological replicates, and the luminescence was normalized by OD_600_ of the yeast co-cultures. The effect of α-factor on the cell growth of R-strains was evaluated using a halo assay as previously reported ^88^. Briefly, yeast strains were grown overnight in 2 ml tubes with 1.7 mL of SC medium at 30ºC with 130 rpm of shaking. Thereafter, 1:10 dilutions of these cultures were merged with 9 mL of soft-SC media (1% agar), and then the mixtures were poured on a plate containing 15 mL of SC media (2% agar). Once solidified, the plates containing R-cells in a uniform lawn were inoculated with 5 µL of commercial pheromone at the same concentrations mentioned above. The solid cultures were grown at 30ºC without shaking for 48 h and pictures of cell plates were taken using the colorimetric mode of Chemidoc Touch Imaging Systems (Bio-Rad, USA). All halo assays were conducted in three biological replicates.

### Evaluation of OS-strains using culture supernatants

BY4741, *bar1Δ* and *bar1Δfar1Δ* strains were co-transformed with the inducible *MFα1* gene construct and the FUN-LOV plasmids, obtaining the Optogenetic Sender (OS) strains. These strains were grown in flasks under darkness overnight in 25 ml of SC medium at 30°C with 130 rpm of shaking. Thereafter, the OD_600_ of the cultures was adjusted to 0.2 and the cells were grown for 8 h under constant darkness (DD) or constant white light (LL) conditions using Percival incubators (Percival Scientific, USA). Importantly, the LL experiments were conducted under 100 µmol m^-2^ s^-1^ of light intensity. Then, the cells were discarded by two consecutive rounds of centrifugation at 3500 rpm for 5 min and the supernatants were recovered and maintained at -20ºC. On the other hand, R-strains were grown overnight in a 96-well plate with 200 µL of SC medium at 30ºC. Thereafter, 10 µL of the R-cell cultures were inoculated in a new 96-well plate containing 190 µL of the collected supernatants supplemented with a final concentration of 1 mM luciferin, to indirectly analyze the optogenetic pheromone production in terms of OD_600_ and luminescence. All kinetic curves were performed with six biological replicates, and the results are shown in terms of luminescence normalized by OD_600_. The growth parameters such as rate and efficiency were calculated using the OD_600_ data and the Gompertz equation ^89^. Also, the pheromone-dependent growth inhibition of R-strains was evaluated with halo assay, using 5 µL of OS-supernatants obtained from DD and LL conditions. Again, the halo assays were conducted in three biological replicates.

### Evaluation of the two-population system

The mixture of OS- and R-strains allowed the evaluation of the optogenetic intercellular system. First, the cells were evaluated at OS-R 1:1 ratio under three different illumination conditions: 24 h of DD, 24 h of blue-light (BL) and a transition of 4 h of DD followed by 20 h of BL (DL4h:20h) in a temperature-controlled dark room. Blue-light (BL) experiments were carried out using an LED lamp (model i5038) at ∼ 20 µmol m^-2^ s^-1^ of light intensity ^50^. Briefly, OS- and R-strains were grown separately overnight in a 96-well plate with 200 µL of SC medium at 30ºC in DD. Thereafter, 5 µL of the OS- and R-cell cultures were co-inoculated in nine combinations in a new 96-well plate containing 190 µL of fresh SC media supplemented with luciferin (1 mM). The measurements involving BL used a discontinuous protocol, that allows keeping the plate outside of the microplate reader to expose the co-cultures to the LED lamp between each measurement ^50^. Then, different dark/blue-light transitions (DL1h:23h, DL2h:22h, DL6h:18h, DL8h:16h, DL10h:14h, DL12h:12h) were assessed. Furthermore, the cells were evaluated by varying the OS-R ratio (3:1 and 1:3) under DD, BL and DL4h:20h conditions. In this case, the normalized results are shown in heat-maps, where each co-culture was evaluated in six biological replicates. Finally, the pheromone-dependent growth inhibition of R-strains was evaluated with halo assay by using 5 µL of saturating cultures of OS-strains, posteriorly grown in DD and LL conditions. As before, these halo assays were conducted in three biological replicates.

## SUPPORTING INFORMATION

The supporting information includes Figure S1 to S15 and Tables S1 to S4.

## AUTHOR INFORMATION

### Author Contributions

Conceptualization, V.R. and L.F.L.; investigation, V.R.; methodology, V.R.; formal analysis, V.R.; visualization, V.R.; draft preparation, V.R. and L.F.L.; review and editing, V.R. and L.F.L.; supervision, L.F.L.; funding acquisition, L.F.L. All authors have read and agreed to the published version of the manuscript.

### Notes

The authors declare no conflict of interest.

## ACKNOWLEDGMENTS

This research was funded by the ANID-Millennium Science Initiative Program-ICN17_022 to L.F.L.; Howard Hughes International Research Scholar program to L.F.L.; ANID-FONDECYT grant number 1211715 to L.F.L.; and by the ANID-Ph.D. scholarship 21170331 to V.R. We thank Dr. Felipe Muñoz-Guzmán for insights and suggestions.

