## Supplementary Figures for "Coupling cell communication and optogenetics: Implementation of a light-inducible intercellular system in yeast"

**SUPPORTING INFORMATION**

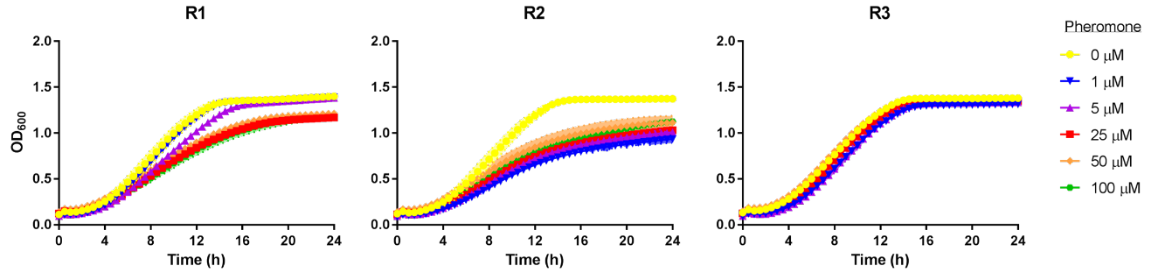


**Figure S1. $OD_{600}$ curves of R-strains supplemented with increasing pheromone concentrations.** Commercial α-factor was exogenously added in different concentrations (1, 5, 25, 50 and 100 μM) and $OD_{600}$ was measured in R1, R2 and R3 cells for 24 h. In all panels, the R-strains were evaluated in the absence of pheromone (0 μM) as negative controls. The average normalized response of six biological replicates is shown, with the standard deviation represented as a region with soft tone.

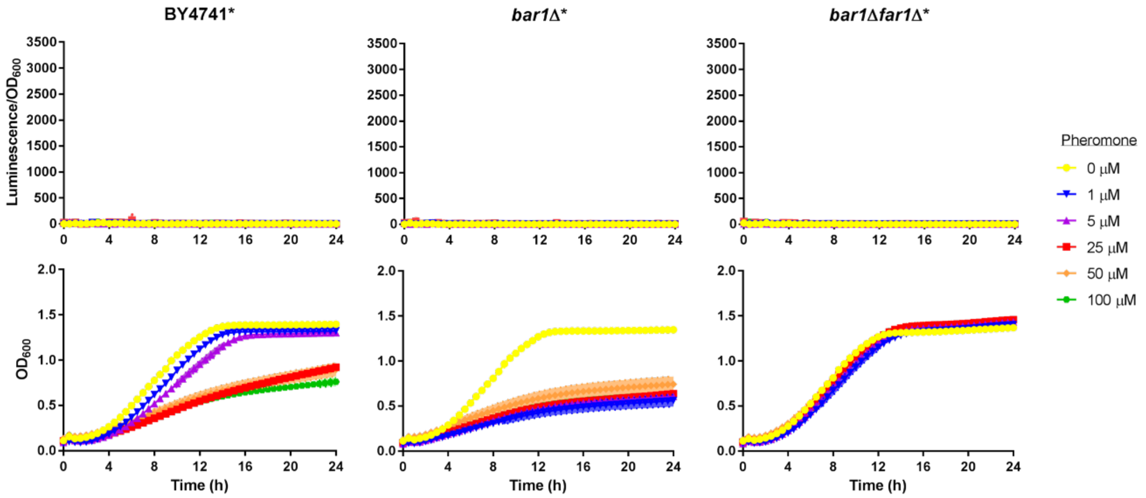


**Figure S2. Normalized response and $OD_{600}$ of strains lacking luciferase gene supplemented with increasing pheromone concentrations.** Commercial α-factor was exogenously added in different concentrations (1, 5, 25, 50 and 100 μM) and normalized reporter expression and $OD_{600}$ were measured in BY4741*, *bar1*Δ* and *bar1*Δ*far1*Δ* cells for 24 h. In all panels, these yeast strains were evaluated in the absence of pheromone (0 μM) as negative controls. The average normalized response of six biological replicates is shown, with the standard deviation represented as a region with soft tone.

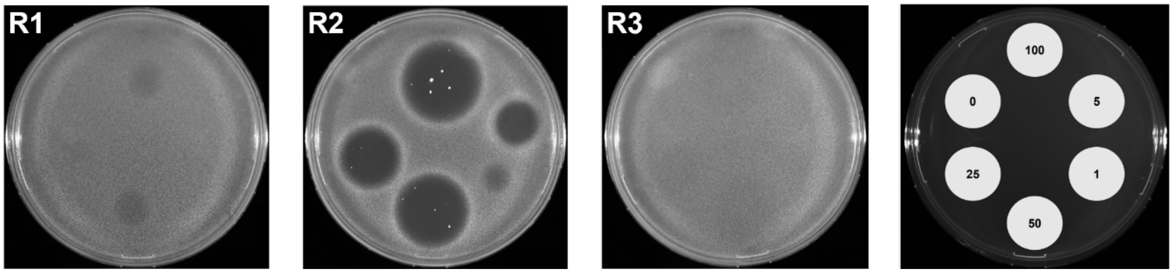


**Figure S3. The R-strains show variable growth alteration upon pheromone addition.** Commercial α-factor was applied in 5 μL drops at different concentrations (1, 5, 25, 50 and 100 μM) over uniform lawns of R1, R2 and R3 cells to qualitatively measure growth inhibition. In all panels, the R-strains were evaluated in the absence of pheromone (0 μM) as negative controls. The plates were cultured by 48 h before taking the pictures. The distribution of concentrations and negative control is shown on the right side of the figure. The images are representatives of three biological replicates for each strain.

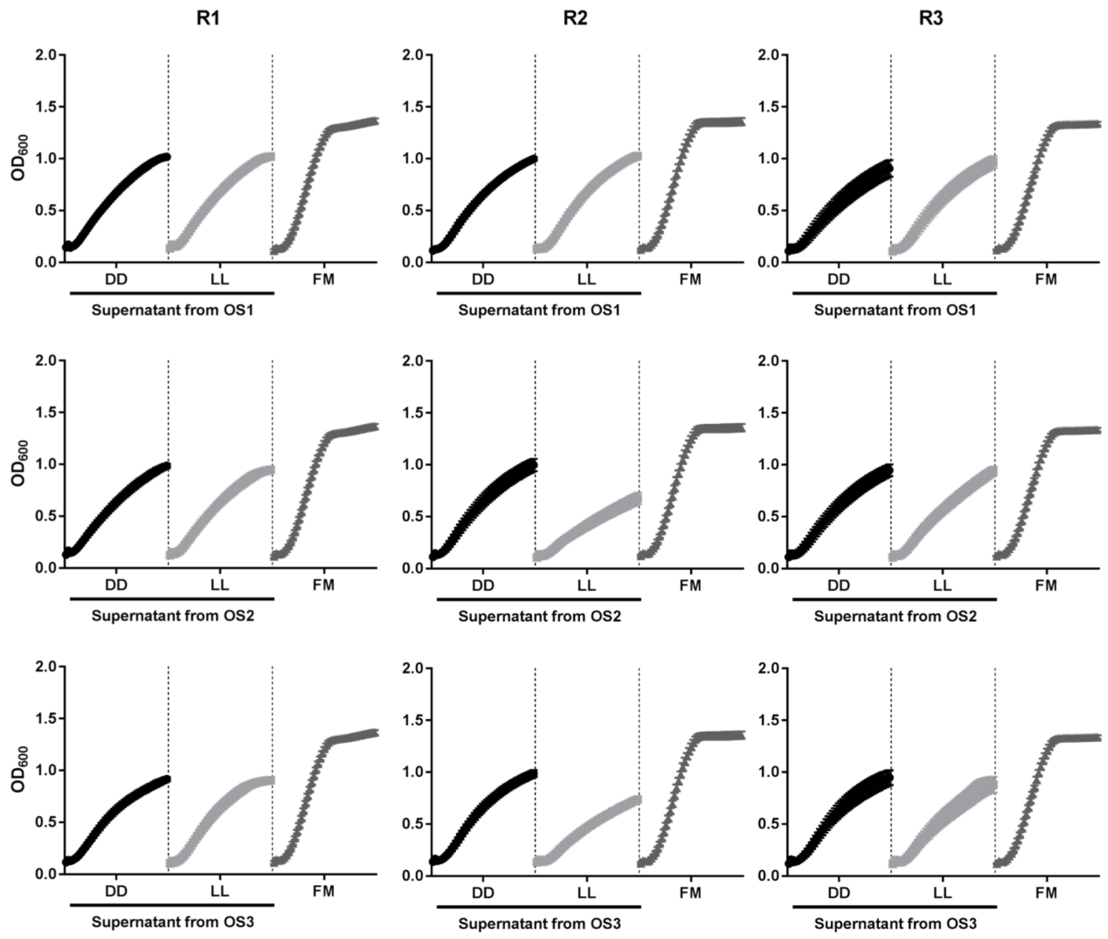


**Figure S4. $OD_{600}$ curves of R-strains grown in supernatants of OS-strains carrying a multi-copy inducible *MFα1* plasmid.** The supernatants of OS1, OS2 and OS3 cultures were collected from contrasting illumination conditions (DD and LL) and $OD_{600}$ was measured in R1, R2 and R3 cells for 24 h. In all panels, the R-strains were evaluated in fresh media (FM) as negative controls. The average normalized response of six biological replicates is shown.

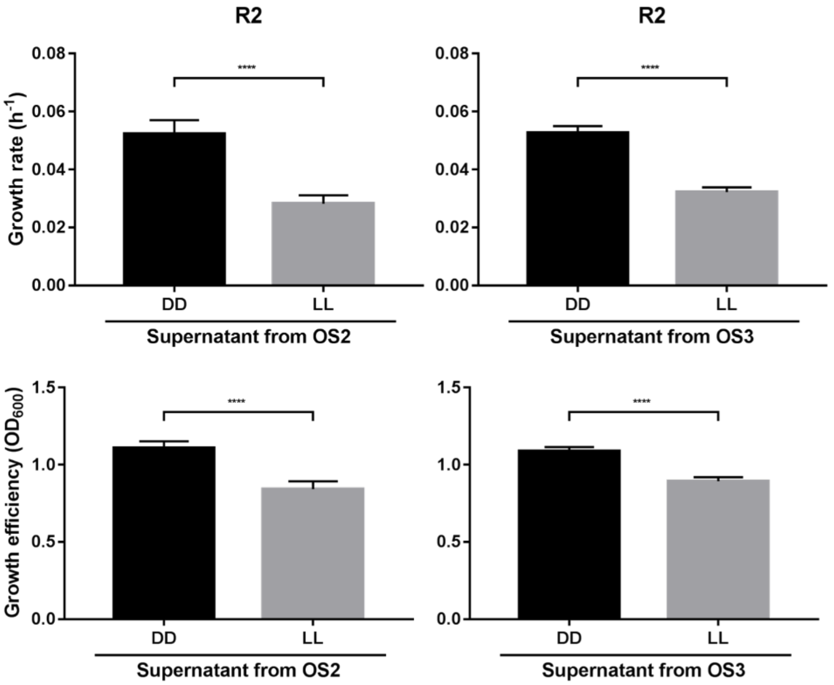


**Figure S5. OS2 and OS3 supernatants obtained in LL affected growth parameters of the R2 strain.** Rate and efficiency were calculated using the $OD_{600}$ curves and Gompertz equation. The average of both parameters in each condition for six biological replicates is shown (**** P <0,0001).

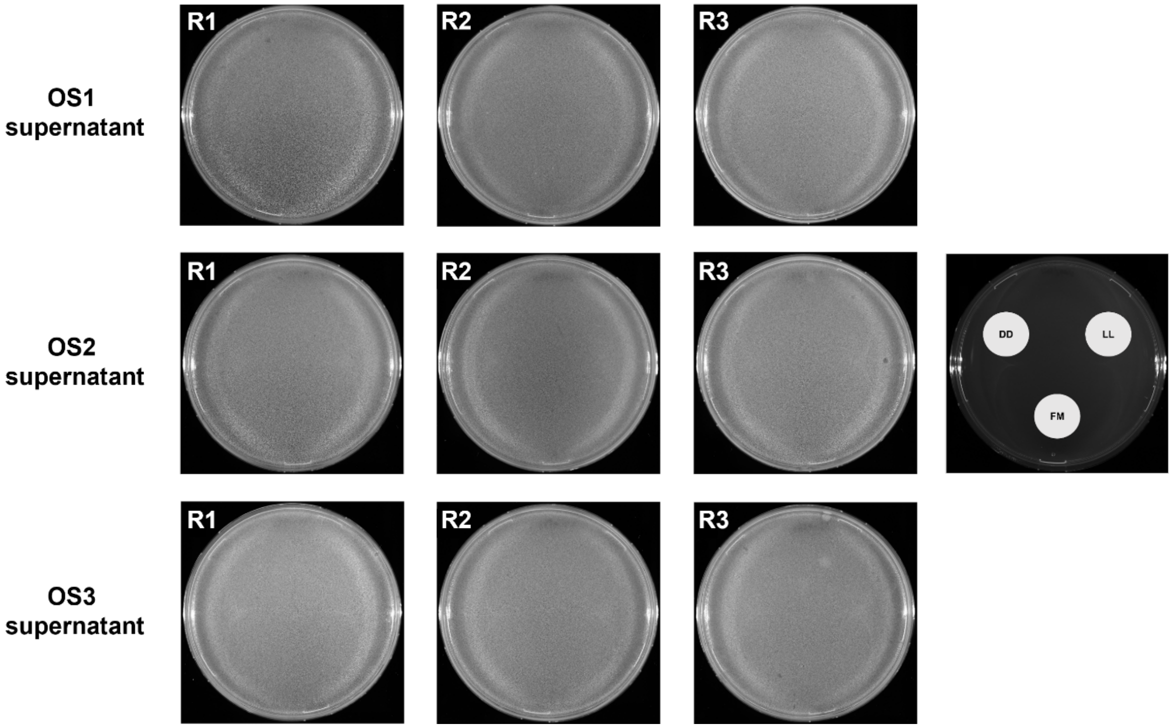


**Figure S6. Halo assays of R-strains using supernatants of OS-strains carrying a multi-copy inducible *MFα1* plasmid.** The supernatants of OS1, OS2 and OS3 cultures were collected from contrasting illumination conditions (DD and LL), and then applied in 5 µL drops over uniform lawns of R1, R2 and R3 cells to qualitatively measure growth inhibition. In all panels, the R-strains were evaluated in fresh media (FM) as negative controls. The plates were cultured by 48 h before taking the pictures. The distribution of supernatants and negative control is shown on the right side of the figure. The images are representatives of three biological replicates for each strain.

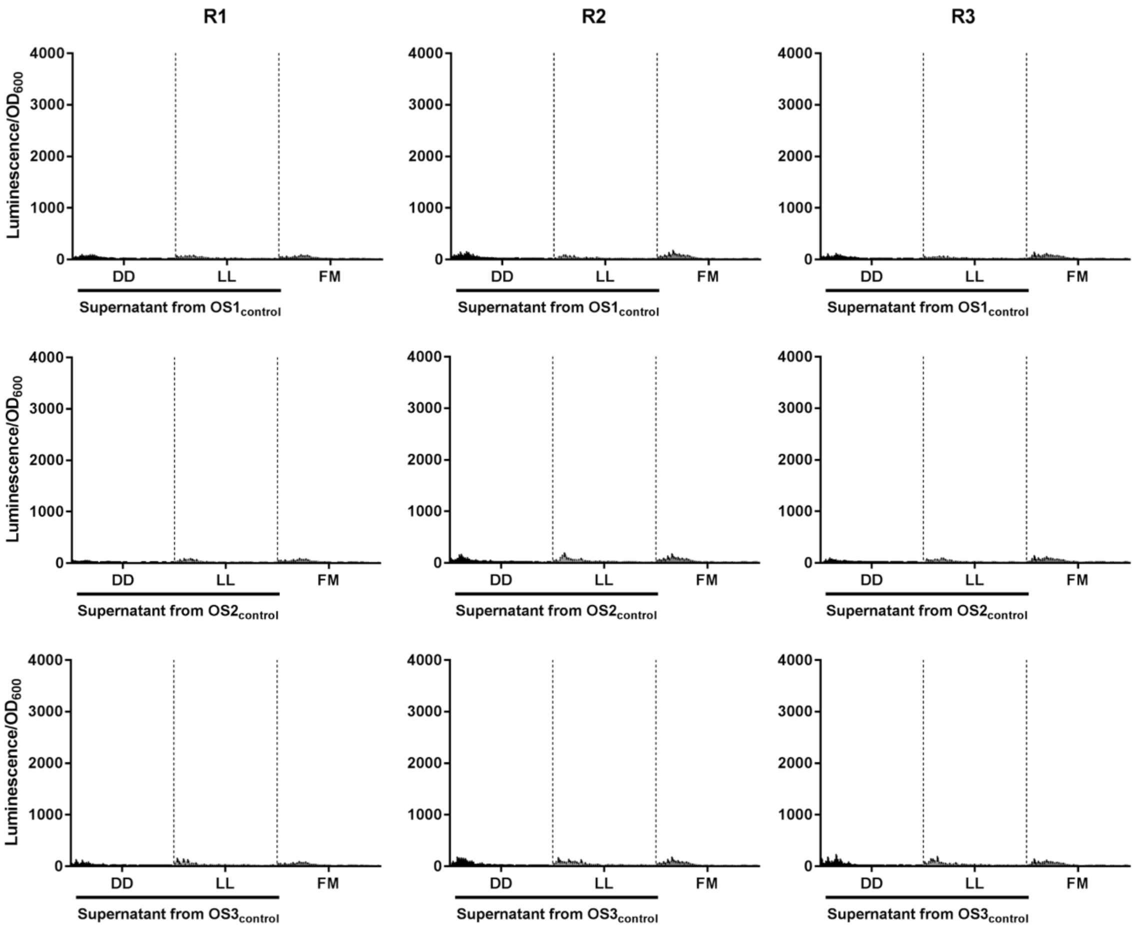


**Figure S7. The OS$_{control}$-strains lacking an inducible *MFα1* construct do not produce functional pheromone upon constant light.** The supernatants of OS1$_{control}$, OS2$_{control}$ and OS3$_{control}$ cultures were collected from contrasting illumination conditions (DD and LL) and reporter expression (luminescence normalized by $OD_{600}$) was measured in R1, R2 and R3 cells for 24 h. In all panels, the R-strains were evaluated in fresh media (FM) as negative controls. The average normalized response of six biological replicates is shown, with the standard deviation represented as a black bar.

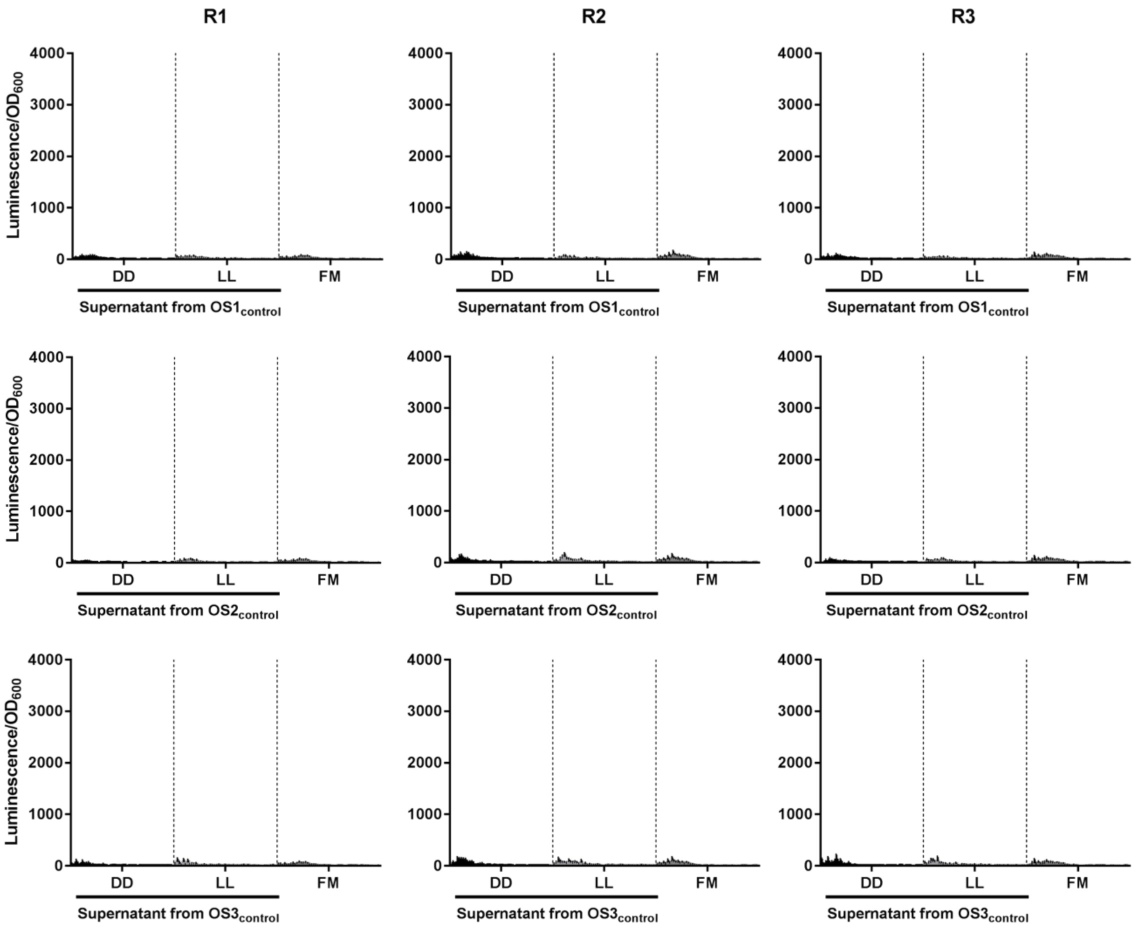


**Figure S8. The OS-strains carrying a chromosomal inducible *MFα1* construct of do not produce functional pheromone upon constant light.** The supernatants of OS4, OS5 and OS6 cultures were collected from contrasting illumination conditions (DD and LL) and reporter expression (luminescence normalized by $OD_{600}$) was measured in R1, R2 and R3 cells for 24 h. In all panels, the R-strains were evaluated in fresh media (FM) as negative controls. The average normalized response of six biological replicates is shown, with the standard deviation represented as a black bar.

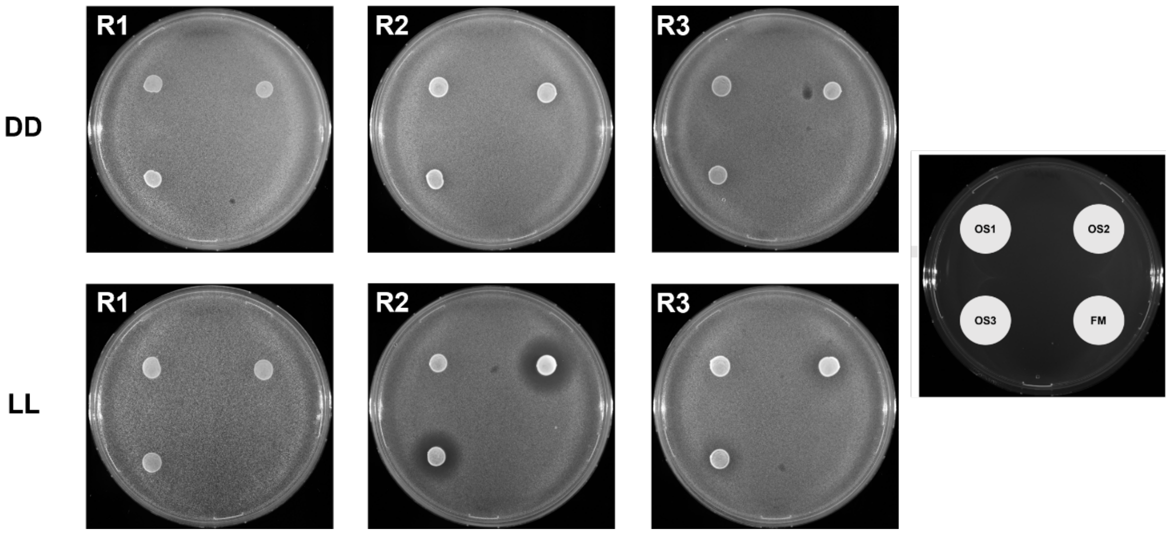


**Figure S9. Halo assays of R-strains using OS-strains carrying a multi-copy inducible *MFα1* plasmid.** The OS1, OS2 and OS3 cells were grown overnight, and then applied in 5 µL drops over uniform lawns of R1, R2 and R3 cells to qualitatively measure growth inhibition. In all panels, the R-strains were evaluated in fresh media (FM) as negative controls. The plates were cultured in DD and LL conditions by 48 h before taking the pictures. The distribution of OS-strains and negative control is shown on the right side of the figure. The images are representatives of three biological replicates for each strain.

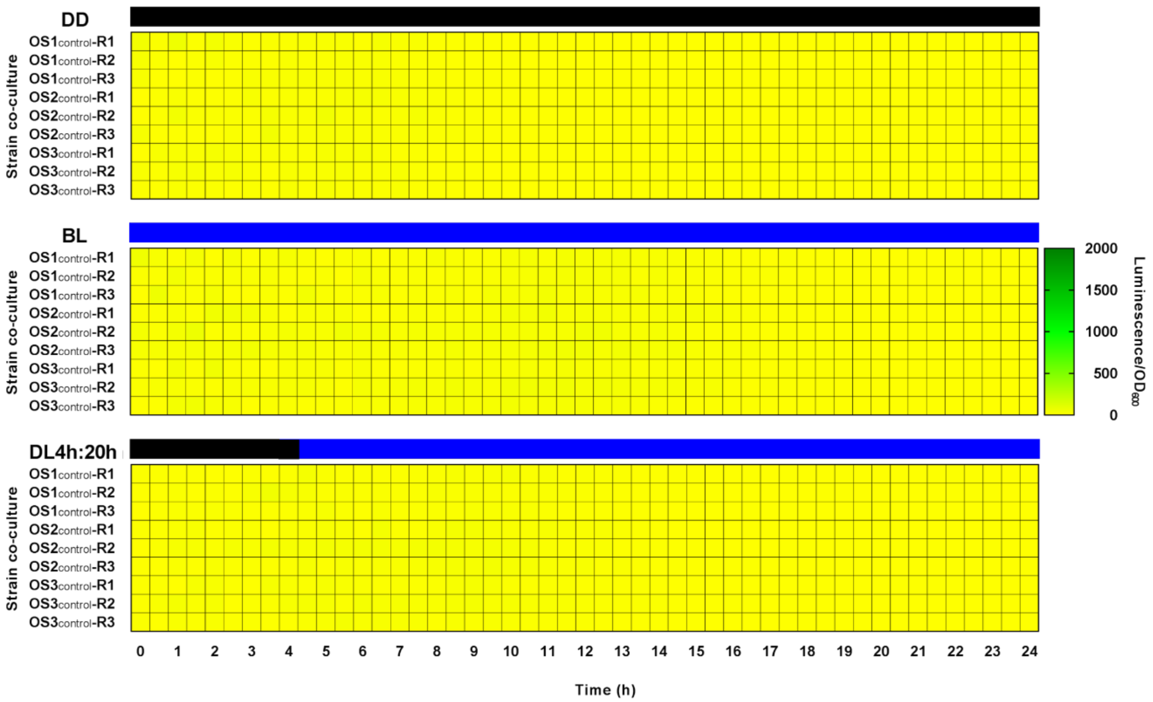


**Figure S10. The combination of $OS_{control}$- and R-strains does not lead to productive light-dependent cell communication.** The cells were combined at $OS_{control}$-R 1:1 ratio, and then exposed to DD, BL and DL4h:20h to measure reporter expression (luminescence normalized by $OD_{600}$) for 24 h. The average normalized response of six biological replicates is shown in the heat-maps.

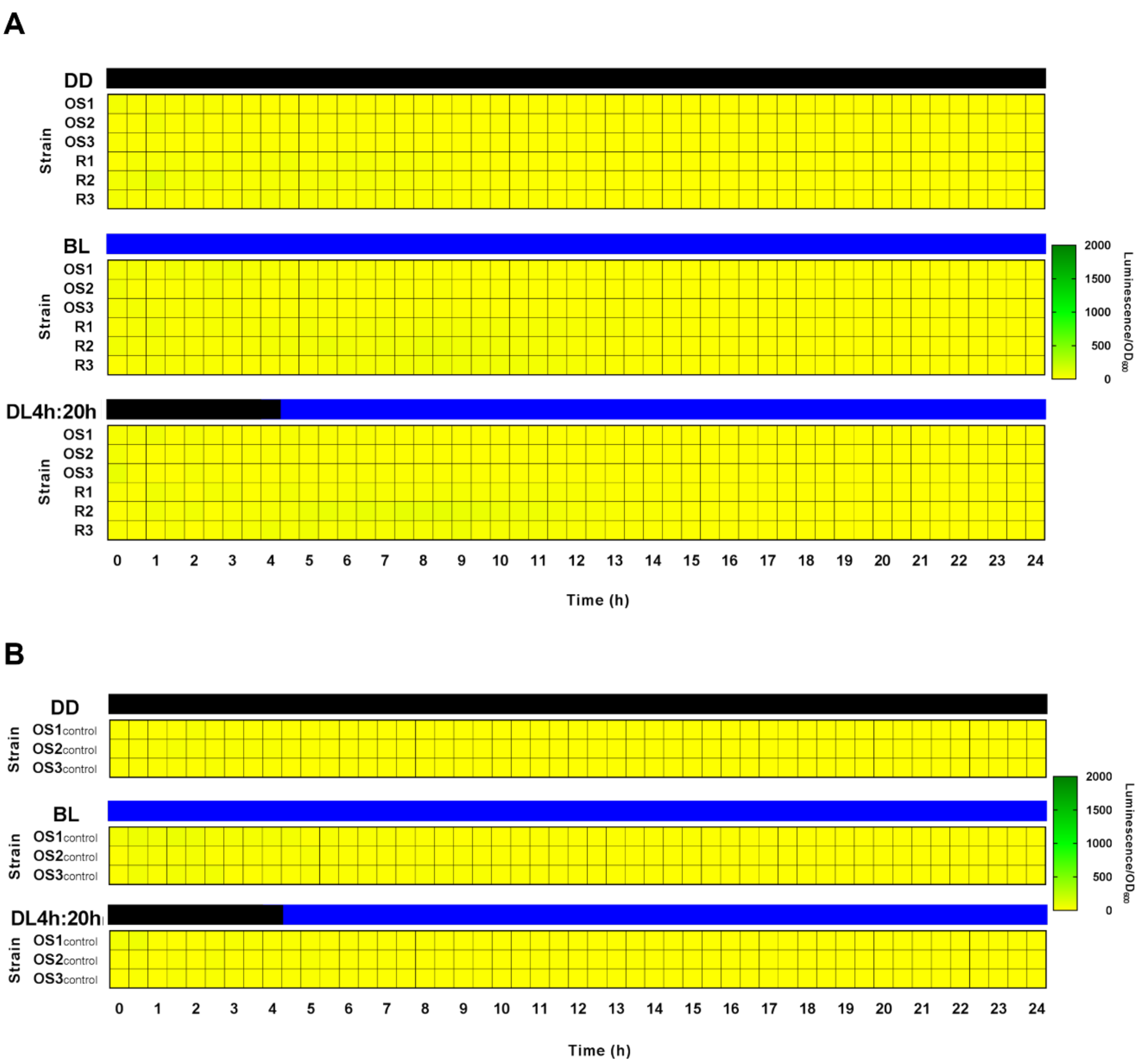


**Figure S11. Normalized response of OS-, R- and $OS_{control}$-monocultures under different illumination conditions.** The cells were cultured individually and then exposed to DD, BL and DL4h:20h to measure reporter expression (luminescence normalized by $OD_{600}$) for 24 h. The average normalized response of six biological replicates is shown in the heat-maps.

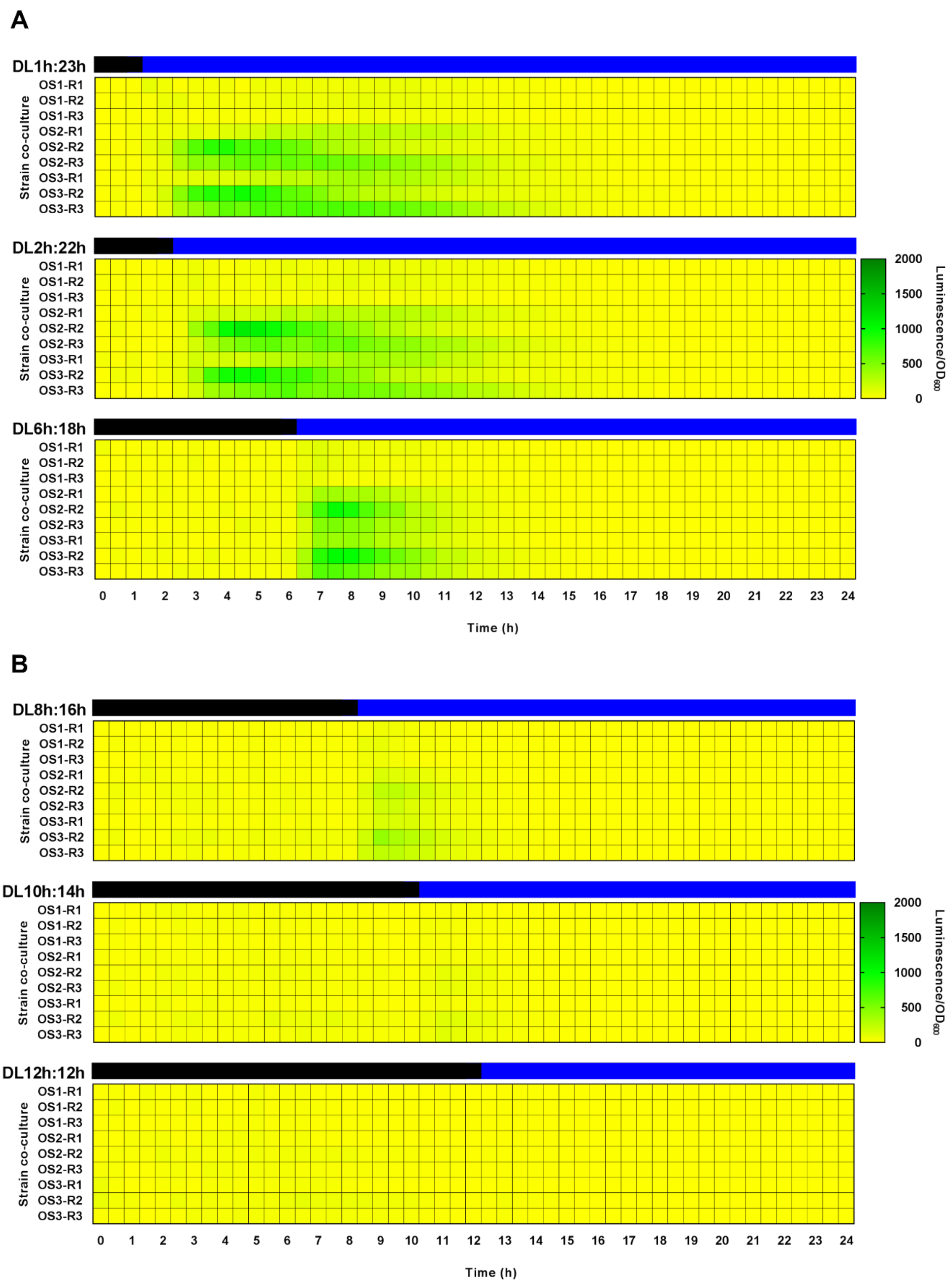


**Figure S12. The timing of illumination is relevant for generating a significant response.** The cells were combined at OS-R 1:1 ratio, and then exposed to early (A) and late (B) dark/blue-light transitions to measure reporter expression (luminescence normalized by $OD_{600}$) for 24 h. The average normalized response of six biological replicates is shown in the heat-maps.

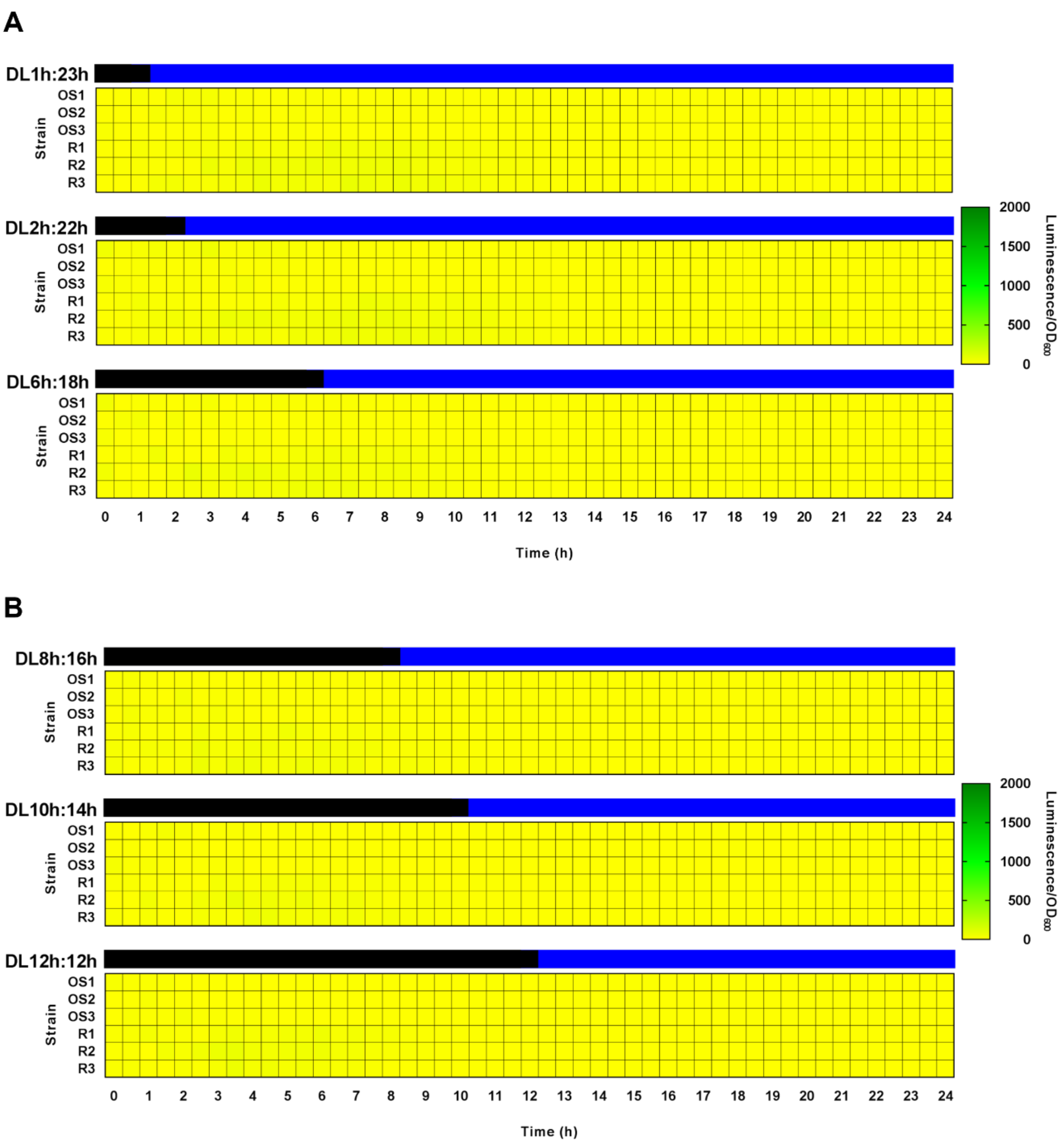


**Figure S13. Normalized response of OS- and R-monocultures under illumination transitions.** The cells were cultured individually and then exposed to early (A) and late (B) dark/blue-light transitions to measure reporter expression (luminescence normalized by $OD_{600}$) for 24 h. The average normalized response of six biological replicates is shown in the heat-maps.

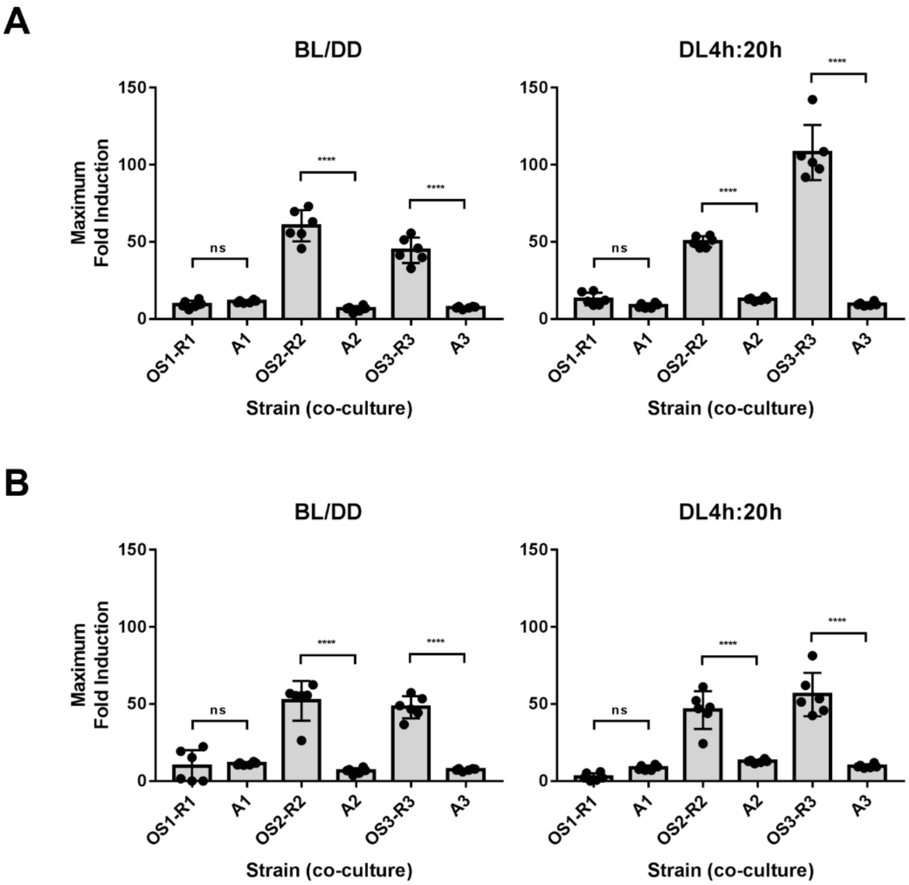


**Figure S14. The autonomous strains display a worse performance regarding asymmetric OS-R co-cultures**. Comparative of maximum fold-induction between A-strains and the optogenetic intercellular system at OS-R 1:3 (A) or 3:1 (B) ratios for two illumination conditions (****P <0,0001).

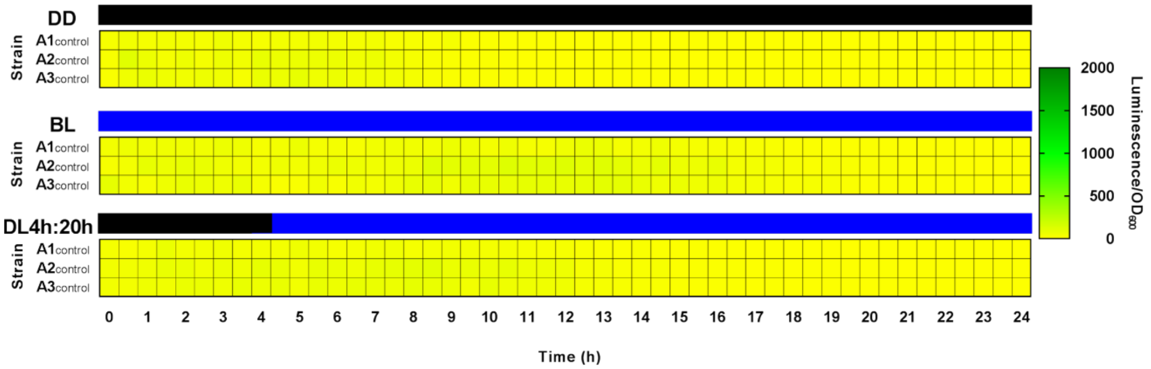


**Figure S15. Normalized response of $A_{control}$-strains under different illumination conditions.** The cells were exposed to DD, BL and DL4h:20h to measure reporter expression (luminescence normalized by $OD_{600}$) for 24 h. The average normalized response of six biological replicates is shown in the heat-maps.

**Table S1. Yeast strains used and generated in this work.**

| Name | Genotype | Source |
|---|---|---|
| BY4741 | *MATa; his3Δ1; leu2Δ0; met15Δ0; ura3Δ0* | Euroscarf |
| *bar1Δ* | BY4741*; bar1Δ::KanMx* | This work |
| *bar1Δfar1Δ* | *bar1Δ; far1Δ::NatMx* | This work |
| BY4741* | BY4741 + pRS423 + pRS425 + pRS426 | This work |
| *bar1Δ** | *bar1Δ* + pRS423 + pRS425 + pRS426 | This work |
| *bar1Δfar1Δ** | *bar1Δfar1Δ* + pRS423 + pRS425 + pRS426 | This work |
| $P_{FUS1}$-*LUC* | BY4741*;* $ORF_{FUS1}$Δ::*LUC-HphMx* | This work |
| *bar1Δ* $P_{FUS1}$-*LUC* | *bar1Δ;* $ORF_{FUS1}$Δ::*LUC-HphMx* | This work |
| *bar1Δfar1Δ* $P_{FUS1}$-*LUC* | *bar1Δfar1Δ;* $ORF_{FUS1}$Δ::*LUC-HphMx* | This work |
| R1 | $P_{FUS1}$-*LUC* + pRS423 + pRS425 + pRS426 | This work |
| R2 | *bar1Δ* $P_{FUS1}$-*LUC* + pRS423 + pRS425 + pRS426 | This work |
| R3 | *bar1Δfar1Δ* $P_{FUS1}$-*LUC* + pRS423 + pRS425 + pRS426 | This work |
| OS1 | BY4741 + WC-1 + VVD + $P_{GAL1}$-*MFα1* | This work |
| OS2 | *bar1Δ* + WC-1 + VVD + $P_{GAL1}$-*MFα1* | This work |
| OS3 | *bar1Δfar1Δ* + WC-1 + VVD + $P_{GAL1}$-*MFα1* | This work |
| $OS1_{control}$ | BY4741 + WC-1 + VVD + pRS426 | This work |
| $OS2_{control}$ | *bar1Δ* + WC-1 + VVD + pRS426 | This work |
| $OS3_{control}$ | *bar1Δfar1Δ* + WC-1 + VVD + pRS426 | This work |
| $P_{GAL1}$-*MFα1* | BY4741*;* $P_{MFα1}$Δ::*URA3*-$P_{GAL1}$ | This work |
| *bar1Δ* $P_{GAL1}$-*MFα1* | *bar1Δ;* $P_{MFα1}$Δ::*URA3*-$P_{GAL1}$ | This work |
| *bar1Δfar1Δ* $P_{GAL1}$-*MFα1* | *bar1Δfar1Δ;* $P_{MFα1}$Δ::*URA3*-$P_{GAL1}$ | This work |
| OS4 | $P_{GAL1}$-*MFα1* + WC-1 + VVD | This work |
| OS5 | *bar1Δ* $P_{GAL1}$-*MFα1* + WC-1 + VVD | This work |
| OS6 | *bar1Δfar1Δ* $P_{GAL1}$-*MFα1* + WC-1 + VVD | This work |
| A1 | $P_{FUS1}$-*LUC* + WC-1 + VVD + $P_{GAL1}$-*MFα1* | This work |
| A2 | *bar1Δ* $P_{FUS1}$-*LUC* + WC-1 + VVD + $P_{GAL1}$-*MFα1* | This work |
| A3 | *bar1Δfar1Δ* $P_{FUS1}$-*LUC* + WC-1 + VVD + $P_{GAL1}$-*MFα1* | This work |
| $A1_{control}$ | $P_{FUS1}$-*LUC* + WC-1 + VVD + pRS426 | This work |
| $A2_{control}$ | *bar1Δ* $P_{FUS1}$-*LUC* + WC-1 + VVD + pRS426 | This work |
| $A3_{control}$ | *bar1Δfar1Δ* $P_{FUS1}$-*LUC* + WC-1 + VVD + pRS426 | This work |

**Table S2. Primers used to generate and check mutant/reporter strains.**

| Name | Sequence (5'-3') | Length (nt) | Type | Description |
|---|---|---|---|---|
| oL3988 | ATCGCCTAAAATCATACCAAAATAAAAAGAGTGTCTA GAAGGGTCATATACGGGTTAATTAAGGCGCGCC | 70 | Fw | *bar1Δ* |
| oL3989 | ACTATATATTTGATATTTATATGCTATAAAGAAATTGT ACTCCAGATTTCATCGATGAATTCGAGCTCGT | 70 | Rv | *bar1Δ* |
| oL3976 | GTCTATAGATCCACTGGAAAGCTTCGTGGGCGTAAG AAGGCAATCTATTACGGGTTAATTAAGGCGCGCC | 70 | Fw | *bar1Δfar1Δ* |
| oL3977 | AAAAAAGGAAAAGCAAAAGCCTCGAAATACGGGCCT CGATTCCCGAACTAATCGATGAATTCGAGCTCGT | 70 | Rv | *bar1Δfar1Δ* |
| oL4867 | TTTTTTCCTTTAAGAGCAGGATATAAGCCATCAAGTT TCTGAAAATCAAAATGGCCGATGCTAAGAACAT | 70 | Fw | $P_{FUS1}$-*LUC* |
| oL3985 | AGTACAGAATTATAGGTATAGATTAAATGCGAACGTC AATATTATTTTCAATCGATGAATTCGAGCTCGT | 70 | Rv | $P_{FUS1}$-*LUC* |
| oL3990 | ATGCGTTGTCCCTGTTTTTC | 20 | Fw | External *BAR1* |
| oL3991 | TTACGGACGTTTAGGATGACG | 21 | Rv | External *BAR1* |
| oL3978 | AACATGCAGCCATTTCACCG | 20 | Fw | External *FAR1* |
| oL3979 | TTACCCGGCTGCGAGTTTT | 19 | Rv | External *FAR1* |
| oL3986 | GTCACTTTTCGCGCTGTCTC | 20 | Fw | External *FUS1* |
| oL3987 | CTGGTGCCGCTCAAATCAAC | 20 | Rv | External *FUS1* |
| oL2091 | CATCCTATGGAACTGCCTCGG | 21 | Fw | Internal *KanMx* |
| oL2090 | TTCAGAAACAACTCTGGCGCA | 21 | Rv | Internal *KanMx* |
| oL2164 | AGGTCACCAACGTCAACGCA | 20 | Fw | Internal *NatMx* |
| oL2163 | GATTCGTCGTCCGATTCGTC | 20 | Rv | Internal *NatMx* |
| oL2095 | TCGCCCGCAGAAGCGCGGCC | 20 | Fw | Internal *HphMx* |
| oL3378 | AGTTCTCAGAGCACACCACG | 20 | Rv | Internal *LUC* |
| oL4658 | CGTATCAATTCCTAAAATCAAAAAAAAAAAAAAAAAAA AGTTAAACAAGCGGTATTTCACACCGCATAGG | 70 | Fw | $P_{GAL1}$-*MFα1* |
| oL4659 | GCGGAGGATGCTGCGAATAAAACTGCAGTAAAAATT GAAGGAAATCTCATGGTAAGCTTAATATTCCCTA | 70 | Rv | $P_{GAL1}$-*MFα1* |
| oL4926 | TCGACATGAAATTTTTGGTATTG | 23 | Fw | External *MFα1* |
| oL4927 | TACATTGGTTGGCCAGGTTT | 20 | Rv | Internal *MFα1* |
| oL2288 | CCTTTTGATGTTAGCAGAATTGTC | 24 | Fw | Internal *URA3* |
| oL3049 | AGTAACCTGGCCCCACAAAC | 20 | Fw | Internal $P_{GAL1}$ |

**Table S3. Recombinant plasmids used and generated in this work.**

| Name | Genetic construct | Vector | Source |
|---|---|---|---|
| WC-1 | $P_{ADH1}$-*WC-1*$^{LOV}$-*GAL4*$^{DBD}$-$T_{ADH2}$ | pRS423 | Salinas et al., 2018 |
| VVD | $P_{ADH1}$-*VVD*$^{LOV}$-*GAL4*$^{AD}$-$T_{ADH2}$ | pRS425 | Salinas et al., 2018 |
| $P_{GAL1}$-*MFα1* | $P_{GAL1}$-*MFα1*-$T_{MF\alpha1}$ | pRS426 | This work |
| *LUC*-HphMx | $P_{ASN1}$-*LUC*$^{PEST}$-$T_{CYC1}$-*HphMx* | pRS426 | Tapia et al., 2018 |
| *URA3*-$P_{GAL1}$ | *URA3*RV-$P_{GAL1}$ | pRS423 | This work |


(1) Salinas, F., Rojas, V., Delgado, V., López, J., Agosin, E., and Larrondo, L. F. (2018) Fungal Light-Oxygen-Voltage Domains for Optogenetic Control of Gene Expression and Flocculation in Yeast. *MBio 9*.

(2) Tapia, S. M., Cuevas, M., Abarca, V., Delgado, V., Rojas, V., García, V., Brice, C., Martínez, C., Salinas, F., Larrondo, L. F., and Cubillos, F. A. (2018) GPD1 and ADH3 natural variants underlie glycerol yield differences in wine fermentation. *Front. Microbiol. 9*, 1–13.

**Table S4. Primers used for plasmid assembly.**

| Name | Sequence (5’-3’) | Length (nt) | Type | Description |
|---|---|---|---|---|
| oL4061 | AGCGGATAACAATTTCACACAGGAAACAGCACTAGTACGGATTAGAAGCC | 50 | Fw | YRP_L-$P_{GAL1}$ |
| oL3980 | AACTGCAGTAAAAATTGAAGGAAATCTCATGGTAAGCTTAATATTCCCTA | 50 | Rv | *MFα1*-$P_{GAL1}$ |
| oL3981 | CGACTCACTATAGGGAATATTAAGCTTACCATGAGATTTCCTTCAATTTTT | 50 | Fw | $P_{GAL1}$-*MFα1* |
| oL3982 | GGTAACGCCAGGGTTTTCCCAGTCACGACGGATTCGATTCACATTCATCT | 50 | Rv | YRP_R-$T_{MF\alpha 1}$ |
| oL3158 | AGCGGATAACAATTTCACACAGGAAACAGCGGTATTTCACACCGCATAGG | 50 | Fw | YRP_L-*URA3RV* |
| oL3161 | CCCGCTCGGCGGCTTCTAATCCGTACTAGTCGGCATCAGAGCAGATTGTA | 50 | Rv | $P_{GAL1}$-*URA3RV* |
| oL3162 | GCACTCTCAGTACAATCTGCTCTGATGCCGACTAGTACGGATTAGAAGCC | 50 | Fw | *URA3RV*- $P_{GAL1}$ |
| oL2528 | GGTAACGCCAGGGTTTTCCCAGTCACGACGGGTAAGCTTAATATTCCCTA | 50 | Rv | YRP_R- $P_{GAL1}$ |
